# Efficient chromatin accessibility mapping *in situ* by nucleosome-tethered tagmentation

**DOI:** 10.1101/2020.04.15.043083

**Authors:** Steven Henikoff, Jorja G. Henikoff, Hatice S. Kaya-Okur, Kami Ahmad

## Abstract

Chromatin accessibility mapping is a powerful approach to identify potential regulatory elements. A popular example is ATAC-seq, whereby Tn5 transposase inserts sequencing adapters into accessible DNA (‘tagmentation’). CUT&Tag is a tagmentation-based epigenomic profiling method in which antibody tethering of Tn5 to a chromatin epitope of interest profiles specific chromatin features in small samples and single cells. Here we show that by simply modifying the tagmentation conditions for histone H3K4me2 or H3K4me3 CUT&Tag, antibody-tethered tagmentation of accessible DNA sites is redirected to produce chromatin accessibility maps that are indistinguishable from the best ATAC-seq maps. Thus, chromatin accessibility maps can be produced in parallel with CUT&Tag maps of other epitopes with all steps from nuclei to amplified sequencing-ready libraries performed in single PCR tubes in the laboratory or on a home workbench. As H3K4 methylation is produced by transcription at promoters and enhancers, our method identifies transcription-coupled accessible regulatory sites.

## Introduction

Identification of DNA accessibility in the chromatin landscape has been used to infer active transcription ever since the seminal description of DNaseI hypersensitivity by Weintraub and Groudine more than 40 years ago (1). Because nucleosomes occupy most of the eukaryotic chromatin landscape and regulatory elements are mostly free of nucleosomes when they are active, DNA accessibility mapping can potentially identify active regulatory elements genome-wide. Several additional strategies have been introduced to identify regulatory elements by DNA accessibility mapping, including digestion with Micrococcal Nuclease (MNase) (2) or restriction enzymes (3), DNA methylation (4), physical fragmentation (5) and transposon insertion (6). With the advent of genome-scale mapping platforms, beginning with microarrays and later short-read DNA sequencing, mapping regulatory elements based on DNaseI hypersensitivity became routine (7, 8). Later innovations included FAIRE (9) and Sono-Seq (10), based on physical fragmentation and differential recovery of cross-linked chromatin, and ATAC-seq (11), based on preferential insertion of the Tn5 transposase. The speed and simplicity of ATAC-seq, in which the cut-and-paste transposition reaction inserts sequencing adapters in the most accessible genomic regions (tagmentation), has led to its widespread adoption in many laboratories for mapping presumed regulatory elements.

For all of these DNA accessibility mapping strategies, it is generally unknown what process is responsible for creating any particular accessible sites within the chromatin landscape. Furthermore accessibility is not all-or-none, with the median difference between an accessible and a non-accessible site in DNA estimated to be only ~20%, with no sites completely accessible or inaccessible in a population of cells (12, 13). Despite these uncertainties, DNA accessibility mapping has successfully predicted the locations of active gene enhancers and promoters genome-wide, with excellent correspondence between methods based on very different strategies (14). This is likely because DNA accessibility mapping strategies rely on the fact that nucleosomes have evolved to repress transcription by blocking sites of preinitiation complex formation and transcription factor binding (15), and so creating and maintaining a nucleosome-depleted region (NDR) is a pre-requisite for promoter and enhancer function.

A popular alternative to DNA accessibility mapping for regulatory element identification is to map nucleosomes that border NDRs, typically by histone marks, including “active” histone modifications, such as H3K4 methylation and H3K27 acetylation, or histone variants incorporated during transcription, such as H2A.Z and H3.3. The rationale for this mapping strategy is that the enzymes that modify histone tails and the chaperones that deposit nucleosome subunits are most active close to the sites of initiation of transcription, which typically occurs bidirectionally at both gene promoters and enhancers to produce stable mRNAs and unstable enhancer RNAs. Although the marks left behind by active transcriptional initiation “point back” to the NDR, this cause-effect connection between the NDR and the histone marks is only by inference (16), and direct evidence is lacking that a histone mark is associated with an NDR.

Here we show that a simple modification of our <u>C</u>leavage <u>U</u>nder <u>T</u>argets& <u>Tagm</u>entation (CUT&Tag) method for antibody-tethered *in situ* tagmentation can identify NDRs genome-wide at regulatory elements adjacent to transcription-associated histone marks in human cells. We provide evidence that reducing the ionic concentration during tagmentation preferentially attracts Tn5 tethered to the H3K4me2 histone modification via a Protein A/G fusion to the nearby NDR, shifting the site of tagmentation from nucleosomes bordering the NDR to the NDR itself. Practically all transcription-coupled accessible sites correspond to ATAC-seq sites and vice-versa, and lie upstream of paused RNA Polymerase II (RNAPII). “CUTAC” (<u>C</u>leavage <u>U</u>nder <u>T</u>argeted <u>A</u>ccessible <u>C</u>hromatin) is conveniently performed in parallel with ordinary CUT&Tag, producing accessible site maps from low cell numbers with signal-to-noise as good as or better than the best ATAC-seq datasets.

## Results

### Streamlined CUT&Tag produces high-quality datasets with low cell numbers

We previously introduced CUT&RUN, a modification of Laemmli’s Chromatin Immunocleavage (ChIC) method (17), in which a fusion protein between Micrococcal Nuclease (MNase) and Protein A (pA-MNase) binds sites of antibodies bound to chromatin fragments in nuclei or permeabilized cells immobilized on magnetic beads. Activation of MNase with Ca^++^ results in targeted cleavage, releasing the antibody-bound fragment into the supernatant for paired-end DNA sequencing. More recently, we substituted the Tn5 transposase for MNase in a modified CUT&RUN protocol, such that addition of Mg^++^ results in a cut-and-paste “tagmentation” reaction, in which sequencing adapters are integrated around sites of antibody binding (18). In CUT&Tag, DNA purification is followed by PCR amplification, eliminating the end-polishing and ligation steps required for sequencing library preparation in CUT&RUN. Like CUT&RUN, CUT&Tag requires relatively little input material, and the low backgrounds permit low sequencing depths to sensitively map chromatin features.

We have developed a streamlined version of CUT&Tag that eliminates tube transfers, so that all steps can be efficiently performed in a single PCR tube (19). However, we had not determined the suitability of the single-tube protocol for profiling low cell number samples. During the COVID-19 pandemic, we adapted this CUT&Tag-direct protocol for implementation with minimal equipment and space requirements that uses no toxic reagents, so that it can be performed conveniently and safely on a home workbench (Figure 1–figure supplement 1). To ascertain the ability of our CUT&Tag-direct protocol to produce DNA sequencing libraries at home with data quality comparable to those produced in the laboratory, we used frozen aliquots of native human K562 cell nuclei prepared in the laboratory and profiled there using the streamlined single-tube protocol. Aliquots of nuclei were thawed and serially diluted in Wash buffer from ~60,000 down to ~60 starting cells, where the average yield of nuclei was ~50%. We used antibodies to H3K4me3, which preferentially marks nucleosomes immediately downstream of active promoters, and H3K27me3, which marks nucleosomes within broad domains of Polycomb-dependent silencing. Aliquots of nuclei were taken home and stored in a kitchen freezer, then thawed and diluted at home and profiled for H3K4me3 and H3K27me3. In both the laboratory and at home we performed all steps in groups of 16 or 32 samples over the course of a single day through the post-PCR clean-up step, treating all samples the same regardless of cell numbers. Whether produced at home or in the lab, all final barcoded sample libraries underwent the same quality control, equimolar pooling, and final SPRI bead clean-up steps in the laboratory prior to DNA sequencing.

Tapestation profiles of libraries produced at home detected nucleosomal ladders down to 200 cells for H3K27me3 and nucleosomal and subnucleosomal fragments down to 2000 cells for H3K4me3 (**Figure 1A-B**). Sequenced fragments were aligned to the human genome using Bowtie2 and tracks were displayed using IGV. Similar results were obtained for both at-home and inlab profiles for both histone modifications (**Figure 1C-D**) using pA-Tn5 produced in the laboratory, and results using commercial Protein A/Protein G-Tn5 (pAG-Tn5) were at least as good. All subsequent experiments reported here were performed at home using commercial pAG-Tn5, which provided results similar to those obtained using batches of homemade pA-Tn5 run in parallel.

**Figure 1.**
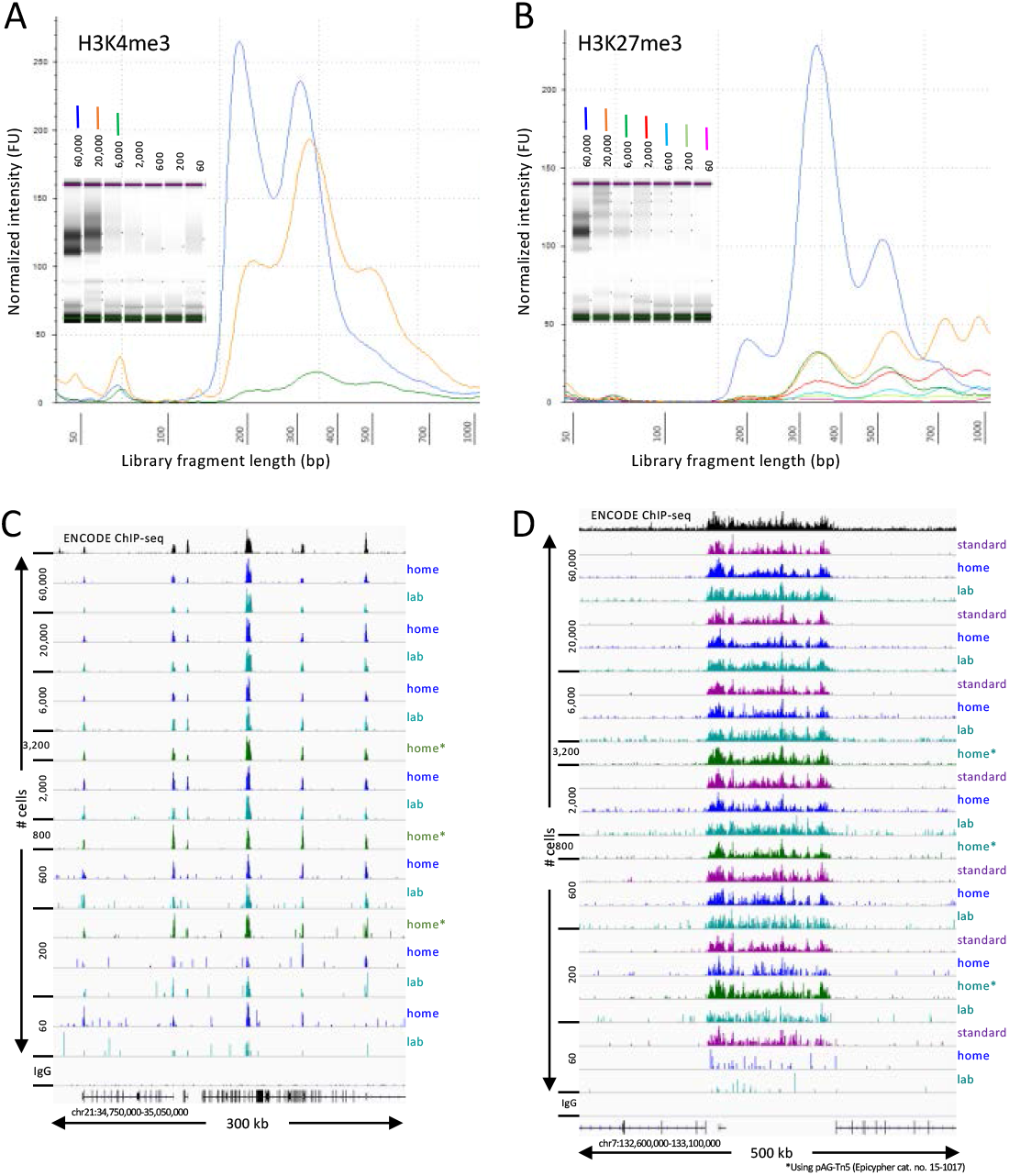
CUT&Tag-direct produces high-quality datasets on the benchtop and at home. Starting with a frozen human K562 cell aliquot, CUT&Tag-direct with amplification for 12 cycles yields detectable nucleosomal ladders for intermediate and low numbers of cells for both (**A**) H3K4me3 and (**B**) H3K27me3. The higher yield of smaller fragments with decreasing cell number suggests that reducing the total available binding sites increases the binding of antibody and/or pAG-Tn5 in limiting amounts. (**C**) Comparison of H3K4me3 CUT&Tag-direct results produced in the laboratory to those produced at home and to an ENCODE dataset (GSM733680). (**D**) Same as (C) for H3K27me3 comparing CUT&Tag-direct results to CUT&Tag datasets using the standard protocol (18), and to an ENCODE dataset (GSM788088). pA-Tn5 was used except as indicated by asterisks for datasets produced at home using commercial pAG-Tn5 (Epicypher cat. no. 15-1017).

### NDRs attract Tn5 tethered to nearby nucleosomes during low-salt tagmentation

Because the Tn5 domain of pA-Tn5 binds avidly to DNA, it is necessary to use elevated salt conditions to avoid tagmenting accessible DNA during CUT&Tag. High-salt buffers included 300 mM NaCl for pA-Tn5 binding, washing to remove excess protein, and tagmentation at 37°C. We have found that other protocols based on the same principle but that do not include a high-salt wash step result in chromatin profiles that are dominated by accessible site tagmentation (19).

To better understand the mechanistic basis for the salt-suppression effect, we bound pAG-Tn5 under normal high-salt CUT&Tag incubation conditions, then tagmented in low salt. We used either rapid 20-fold dilution with a prewarmed solution of 2 mM or 5 mM MgCl2 or removal of the pAG-Tn5 incubation solution and addition of 50 μL 10 mM TAPS pH8.5, 5 mM MgCl2. All other steps in the protocol followed our CUT&Tag-direct protocol (19) (**Figure 2)**. Tapestation capillary gel electrophoresis of the final libraries revealed that after a 20 minute incubation the effect of low-salt tagmentation on H3K4me2 CUT&Tag samples was a marked reduction in the oligo-nucleosome ladder with an increase in faster migrating fragments (**Figure 3A and Figure 3–figure supplement 1A-B**). CUT&Tag profiles using antibodies to most chromatin epitopes in the dilution protocol showed either little change or elevated levels of non-specific background tagmentation that obscured the targeted signal (**Figure 3–figure supplement 2**), as expected considering that we had omitted the high-salt wash step needed to remove unbound pAG-Tn5. Strikingly, under low-salt conditions, high-resolution profiles of H3K4me3 and H3K4me2 showed that the broad nucleosomal distribution of CUT&Tag around promoters for these two modifications was mostly replaced by single narrow peaks (**Figure 3B and Figure 3–figure supplement 3**).

**Figure 2.**
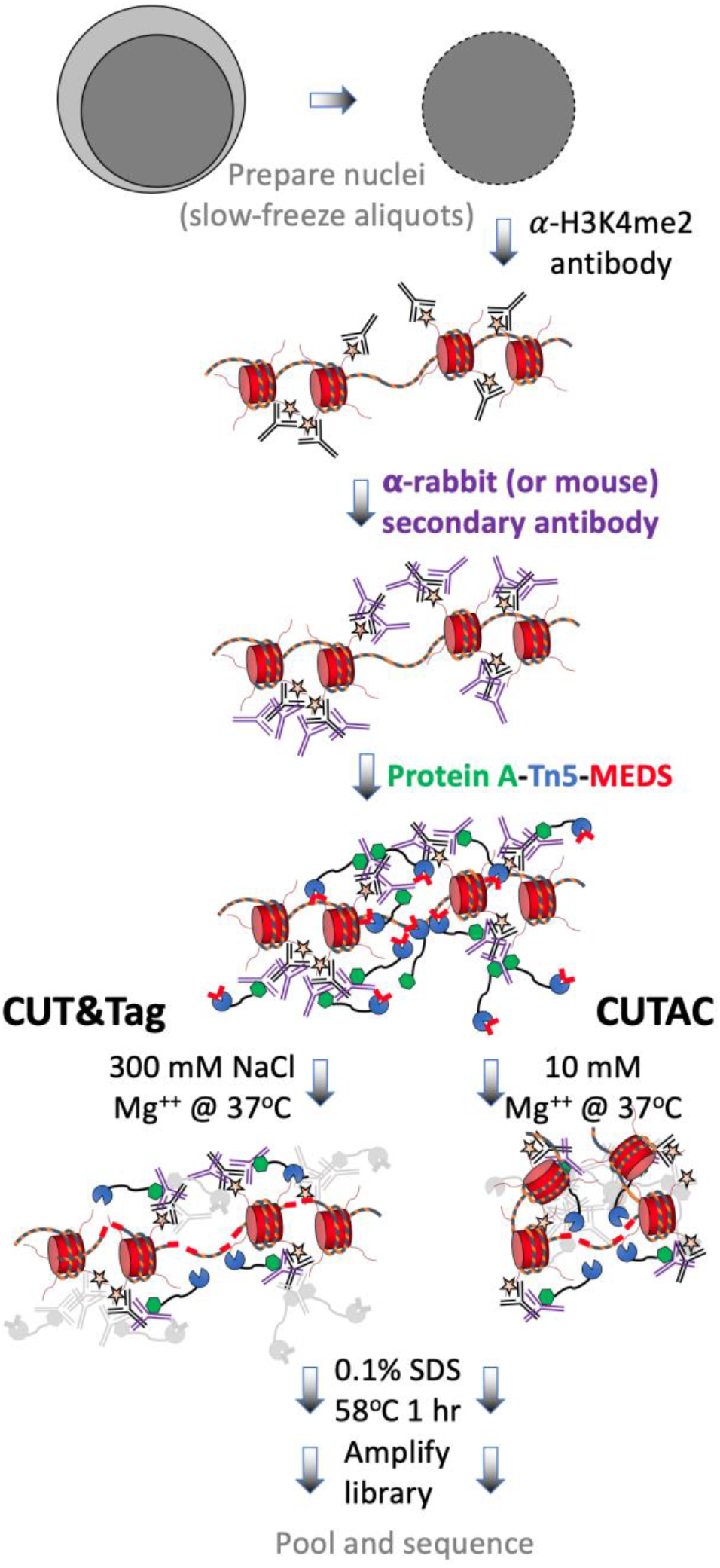
CUT&Tag with low-salt tagmentation (CUTAC). Steps in grey are lab-based and other steps were performed at home. Tagmentation can be performed by dilution, removal or post-wash. MEDS (Mosaic End Double-Stranded annealed oligonucleotides).

**Figure 3.**
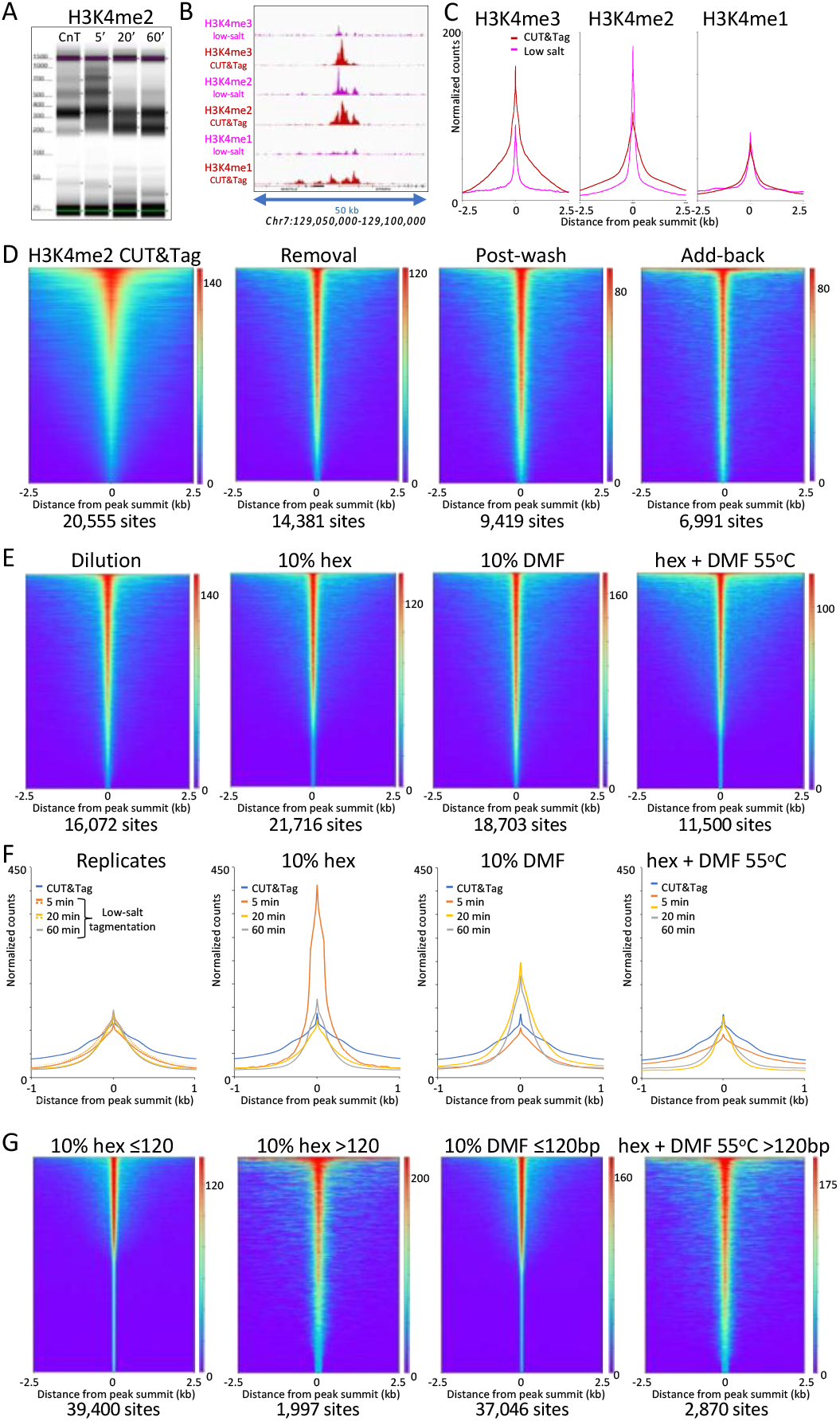
Low-salt tagmentation of H3K4me2/3 CUT&Tag samples sharpen peaks. (**A**) Tapestation gel image showing the change in size distribution from standard CUT&Tag (CnT), tagmented in the presence of 300 mM NaCl with low-salt tagmentation using the dilution protocol. (**B**) Representative tracks showing the shift observed with low-salt dilution tagmentation. (**C**) Average plots showing the narrowing of peak distributions upon low-salt tagmentation using the dilution protocol. (**D**) Heatmaps showing narrowing of H3K4me2 peaks after removing pAG-Tn5 (Removal), after a stringent wash (Post-wash), and after a stringent wash with low-salt tagmentation including a 1% pAG-Tn5 spike-in (Add-back). MACS2 was used to call peaks and heatmaps were ordered by density over the peak midpoints (sites). (**E**) Heatmaps showing dilution tagmentation and further narrowing of H3K4me2 peak distributions upon low-salt tagmentation (after removal) for 20 minutes at 37°C in the presence of 10% 1,6- hexanediol (hex) and 10% dimethylformamide (DMF) or both for 1 hr at 55°C. (**F**) Average plots showing effects of tagmentation with hex and/or DMF over time of low-salt tagmentation (after removal). (**G**) Smaller fragments (≤120 bp) dominate NDRs. Comparisons of small (≤120 bp) and large (>120) fragments from CUTAC hex and DMF datasets show narrowing for small fragments around their summits. For each dataset a 3.2 million fragment random sample was split into small and large fragment groups, Removal of large fragments increases number of peaks called (sites).

To evaluate the generality of peak shifts we used MACS2 to call peaks, and plotted the occupancy over aligned peak summits. For all three H3K4 methylation marks using normal CUT&Tag high-salt tagmentation conditions we observed a bulge around the summit representing the contribution from adjacent nucleosomes on one side or the other of the peak summit (**Figure 3C)**. In contrast, tagmentation under low-salt conditions revealed much narrower profiles for H3K4me3 and H3K4me2 (~40% peak width at half-height), less so for H3K4me1 (~60%), which suggests that the shift is from H3K4me-marked nucleosomes to an adjacent NDR.

To determine whether free pAG-Tn5 present during tagmentation contributes, we removed the pAG-Tn5 then added 5 mM MgCl2 to tagment, and again observed narrowing of the H3K4me2 peak (**Figure 3D “Removal” and Figure 3–figure supplement 3C-D)**. We also observed a narrowing if we included a stringent 300 mM washing step before low-salt tagmentation (**Figure 3D, “Post-wash”**), which indicates that peak narrowing does not require free pAG-Tn5. Inclusion of a stringent post-wash step improves consistency relative to the Dilution or Removal protocols, although it resulted in lower yields and reduced library complexity (Figure 3–figure supplement 3E-F). However, if a small amount of pAG-Tn5 was included during tagmentation we obtained higher yields with increased peak narrowing (**Figure 3D “Add-back”**). Because Tn5 is inactive once it integrates its payload of adapters, and each fragment is generated by tagmentation at both ends, it is likely that a small amount of free pA(G)-Tn5 is sufficient to generate the additional small fragments where tethered pA(G)-Tn5 is limiting, albeit with higher background.

Salt ions compete with protein-DNA binding and so we suppose that tagmentation in low salt resulted in increased binding of epitope-tethered Tn5 to a nearby NDR prior to tagmentation. As H3K4 methylation is deposited in a gradient of tri- to di- to monomethylation downstream of the +1 nucleosome from the transcriptional start site (TSS) (20, 21), we reasoned that the closer proximity of di- and tri-methylated nucleosomes to the NDR than mono-methylated nucleosomes resulted in preferential proximity-dependent “capture” of Tn5. Consistent with this interpretation, we observed that the shift from broad to more peaky NDR profiles and heatmaps by H3K4me2 low-salt tagmentation was enhanced by addition of 1,6-hexanediol, a strongly polar aliphatic alcohol, and by 10% dimethylformamide, a strongly polar amide, both of which enhance chromatin accessibility (**Figure 3E-F**). NDR-focused tagmentation persisted even in the presence of both strongly polar compounds at 55°C. Enhanced localization by chromatin-disrupting conditions suggests improved access of H3K4me2-tethered Tn5 to nearby holes in the chromatin landscape during low-salt tagmentation. Localization to NDRs is more precise for small (≤120 bp) than large (>120) tagmented fragments, and by resolving more closely spaced peaks inclusion of these compounds increased the number of peaks called (**Figure 3D**), also for H3K4me3-tethered Tn5 (**Figure 3–figure supplement 4**).

### CUT&Tag low-salt tagmentation fragments coincide with ATAC-seq and DNaseI hypersensitive sites

Using CUT&Tag, we previously showed that most ATAC-seq sites are flanked by H3K4me2-marked nucleosomes in K562 cells (18). However, lining up ATAC-seq datasets over peaks called using H3K4me2 CUT&Tag data resulted in smeary heatmaps, reflecting the broad distribution of peak calls over nucleosome positions flanking NDRs (**Figure 4A**). In contrast, alignment of ATAC-seq datasets over peaks called using low-salt tagmented CUT&Tag data produced narrow heatmap patterns for the vast majority of peaks (**Figure 4B**). To reflect the close similarities between fragments released by H3K4me2-tethered low-salt tagmentation as by ATAC-seq using untethered Tn5, we will refer to low-salt H3K4me2 and H3K4me3 CUT&Tag tagmentation as <u>C</u>leavage <u>U</u>nder <u>T</u>argeted <u>A</u>ccessible <u>C</u>hromatin (CUTAC).

**Figure 4.**
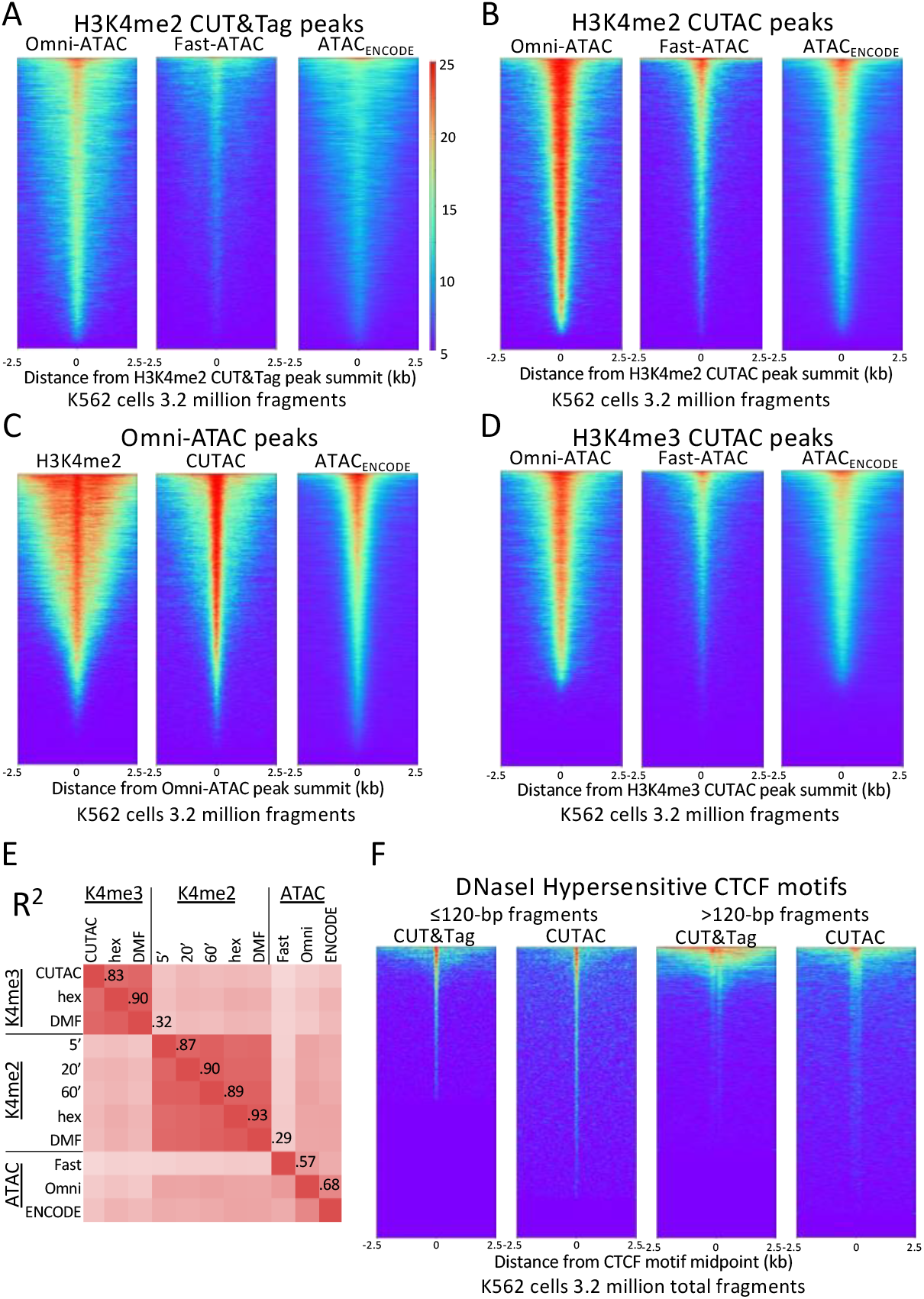
H3K4me2 CUTAC sites coincide with ATAC-seq and DNaseI hypersensitive sites. (**A-D**) Heatmaps showing the correspondence between H3K4me2 CUTAC and ATAC-seq sites. Headings over each heatmap denote the source of fragments mapping to the indicated set of MACS2 peak summits, ordered by occupancy over the 5-kb interval centered over each site. CUT&Tag and CUTAC sites are from samples processed in parallel, where CUTAC tagmentation was performed by 20-fold dilution and 20 minute 37°C incubation following pAG-Tn5 binding. (**E**) Correlation matrix of H3K4me2 and H3K4me3 CUTAC and ATAC-seq data for K562 cells. (**F**) Heatmaps showing that ~90% of CTCF DNaseI hypersensitive sites are detected by H3K4me2 CUTAC.

We confirmed the similarity between CUTAC and ATAC-seq by aligning H3K4me2 CUT&Tag and CUTAC datasets over peaks called from Omni-ATAC data (**Figure 4C**). In a scatterplot comparison between CUTAC and Omni-ATAC we did not detect off-diagonal clusters that would indicate a subset of peaks found by one but not the other dataset (**Figure 4–figure supplement 1**).

To further evaluate the degree of similarity between CUTAC and ATAC-seq, we aligned the ENCODE ATAC-seq dataset over peaks called using Omni-ATAC and CUTAC, where all datasets were sampled down to 3.2 million mapped fragments with mitochondrial fragments removed. Remarkably, heatmaps produced using either Omni-ATAC or CUTAC peak calls for the same ENCODE ATAC-seq data showed occupancy of ~95% for both sets of peaks (compare right panels of **Figure 4B-C**). Using a window of 250 bp around the peak summit based on average peak width at half-height, we found ~50% overlap between ENCODE ATAC-seq peaks and peaks called from either Omni-ATAC (50.0%) or CUTAC (51.3%) data. This equivalence between H3K4me2 CUTAC and Omni-ATAC when compared to ENCODE ATAC-seq implies that CUTAC and Omni-ATAC are indistinguishable in detecting the same chromatin features. This conclusion does not hold for H3K4me3 CUTAC, because similar alignment of ENCODE ATAC-seq data resulted in only ~75% peak occupancy (**Figure 4D**) and lower correlations (**Figure 4E**), which we attribute to the greater enrichment of H3K4me3 around promoters than enhancers relative to H3K4me2.

To evaluate whether CUTAC peaks also correspond to sites of DNaseI hypersensitivity, we aligned H3K4me2 CUT&Tag and CUTAC signals over 9403 CCCTC-binding factor (CTCF) motifs scored as peaks of DNaseI sensitivity in K562 and HeLa cells. We excluded nucleosomal fragments by using only ≤120 bp fragments. We observed that >90% of the DNaseI hypersensitive CTCF sites are occupied by CUTAC signal relative to flanking regions (Figure 4F), which suggests equivalence of CUTAC and DNaseI hypersensitive CTCF sites. We also found that the H3K4me2 CUT&Tag sample showed detectable signal at only ~50% of the CTCF sites. This improvement in detection of CTCF sites by H3K4me2 CUTAC over H3K4me2 CUT&Tag illustrates the potential of using ≤120-bp CUTAC fragment data to improve the resolution and sensitivity of transcription factor binding site motif detection.

To evaluate signal-to-noise genome-wide, we called peaks using MACS2 and calculated the Fraction of Reads in Peaks (FRiP), a data quality metric introduced by the ENCODE project (22). For both ENCODE ChIP-seq and our published CUT&RUN data we measured FRiP ~ 0.2 for 3.2 million fragments, whereas for CUT&Tag, FRiP ~ 0.4, reflecting improved signal-to-noise relative to previous chromatin profiling methods (18). Using CUT&Tag-direct, H3K4me2 CUT&Tag FRiP = 0.41 for 3.2 million fragments and ~16,000 peaks (n=4), whereas tagmentation by dilution in 2 mM MgCl2 resulted in FRiP = 0.18 for 3.2 million fragments and ~15,000 peaks (n=4) with similar values for tagmentation by removal [FRiP = 0.21, ~15,000 peaks (n=4)]. In add-back experiments, we measured lower FRiP values after stringent washing conditions, suggesting increased background.

We also compared the number of peaks and FRiP values for CUTAC to those for ATAC-seq for K562 cells and observed that CUTAC data quality was similar to that for the Omni-ATAC method (23), better than ENCODE ATAC-seq (24), and much better than Fast-ATAC (25), a previous improvement over Standard ATAC-seq (11) (**Figure 5A**). CUTAC is relatively insensitive to tagmentation times, with similar numbers of peaks and similar FRiP values for samples tagmented for 5, 20 and 60 minutes (**Figure 5A**). We attribute the robustness of CUT&Tag and CUTAC to the tethering of Tn5 to specific chromatin epitopes, so that when tagmentation goes to completion there is little untethered Tn5 that would increase background levels. When we measured peak numbers and FRiP values for ATAC-seq for K562 data deposited in the Gene Expression Omnibus (GEO) from multiple laboratories, we observed a wide range of data quality (**Figure 5B**, even from very recent submissions from expert groups (**Table 1 and Figure 5–figure supplement 1**). We attribute this variability to the difficulty of avoiding background tagmention by excess free Tn5 in ATAC-seq protocols and subsequent release of non-specific nucleosomal fragments (26).

**Figure 5.**
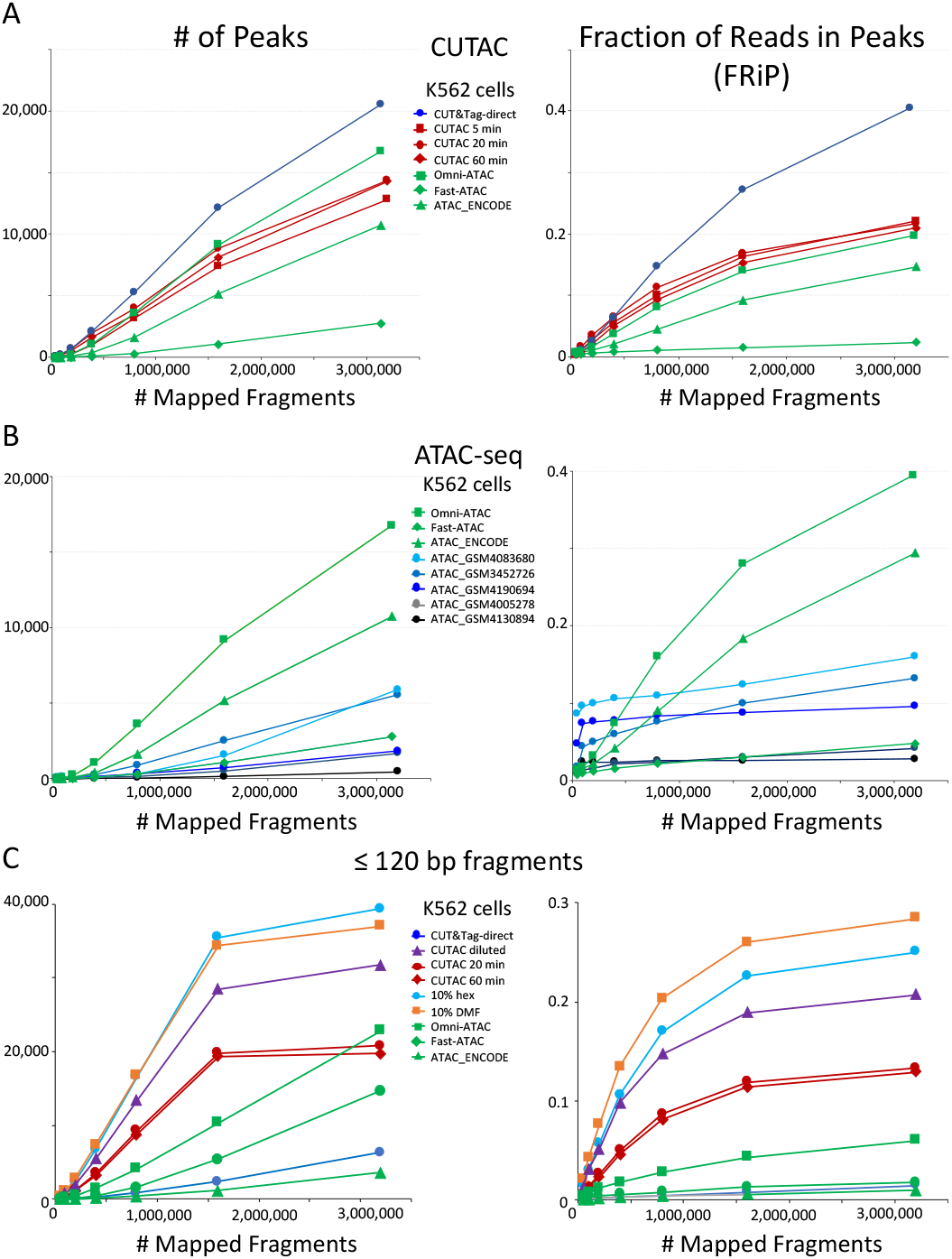
CUTAC data quality is similar to the best available ATAC-seq K562 cell data. Mapped fragments from the indicated datasets were sampled and mapped to hg19 using Bowtie2, and peaks were called using MACS2. (**A**) Number of peaks (left) and fraction of reads in peaks for CUT&Tag (blue), H3K4me2 CUTAC (red) and ATAC-seq (green). Fast-ATAC is an improved version of ATAC-seq that reduces mitochondrial reads (25), and Omni-ATAC is an improved version that additionally improves signal-to-noise (23). ATAC_ENCODE is the current ENCODE standard (35). (**B**) Five other K562 ATAC-seq datasets from different laboratories were identified in GEO and mapped to hg19. MACS2 was used to call peaks and peak numbers, and FRiP values indicate a wide range of data quality found in recent ATAC-seq datasets. (**C**) Small H3K4me2 CUTAC fragments improve peak-calling. Hex = 1,6- hexanediol, DMF = N,N-dimethylformamide.

**Table 1.**
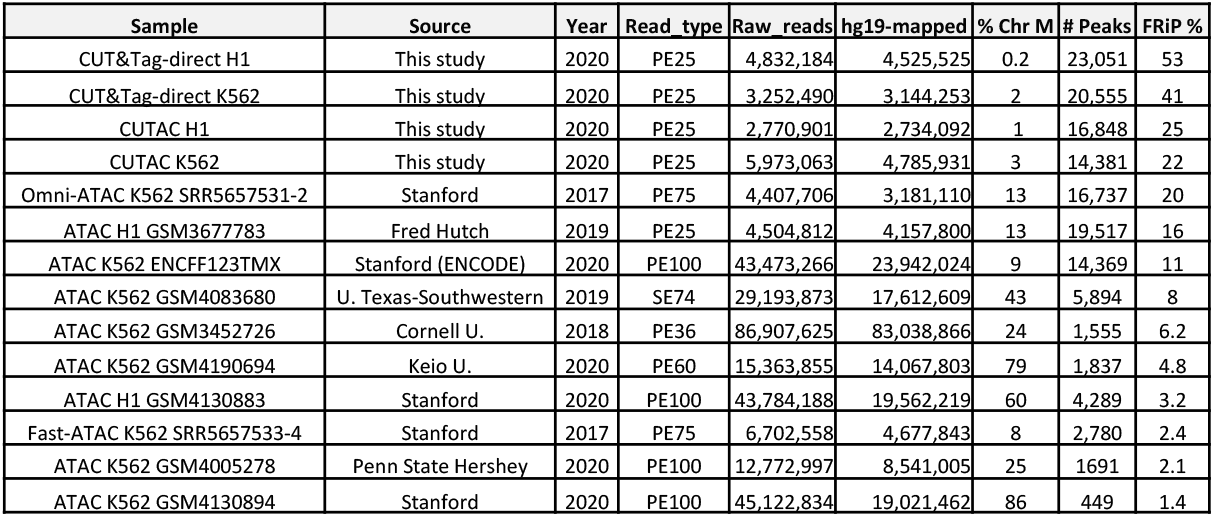
CUTAC data quality is similar to that of the best ATAC-seq datasets. Human K562 and H1 ES cell ATAC-seq datasets were downloaded from GEO, and Bowtie2 was used to map fragments to hg19. A sample of 3.2 million mapped fragments without Chr M was used for peak-calling by MACS2 to calculate FRiP values. Year of submission to GEO or SRA databanks is shown. % Chr M is percent of fragments mapped to Chr M (mitochondrial DNA).

If low-salt tagmentation sharpens peaks of DNA accessibility because tethering to neighboring nucleosomes increases the probability of tagmentation in small holes in the chromatin landscape, then we would expect smaller fragments to dominate CUTAC peaks. Indeed this is exactly what we observe for heatmaps (**Figure 5–figure supplement 2**), tracks (**Figure 5– figure supplement 3**), peak calls and FRiP values (**Figure 5C**). Excluding larger fragments results in better resolution yielding more peaks and higher FRIP values, both of which approach a maximum with fewer fragments. Moreover, the addition of strongly polar compounds during tagmentation provides a substantial improvement in peak calling and FRiPs (**Figure 5C**, turquoise and orange curves). Excluding large fragments did not improve ATAC-seq peak calls and FRiP values, which indicates that tethering to H3K4me2 is critical for maximum sensitivity and resolution of DNA accessibility maps.

### CUTAC maps transcription-coupled regulatory elements

H3K4me2/3 methylation marks active transcription at promoters (27), which raises the question as to whether sites identified by CUTAC are also sites of RNAPII enrichment genome-wide. To test this possibility, we first aligned CUT&Tag and CUTAC data at annotated promoters displayed as heatmaps or average plots. CUT&Tag H3K4me2 peaks flank NDRs more downstream on either side than H3K4me3, confirmed by ENCODE ChIP-seq data to be the actual location of these marks (**Figure 6–figure supplement 1**). In contrast, CUTAC peaks are located in the NDR between flanking H3K4me2-marked chromatin (**Figure 6A**). CUTAC sites at promoter NDRs corresponded closely to promoter ATAC-seq sites, consistent with expectation for promoter NDRs. Thus, paired CUT&Tag and CUTAC samples can replace both ChIP-seq for an active promoter mark and ATAC-seq in a single experiment with identical processing, analysis and display.

**Figure 6.**
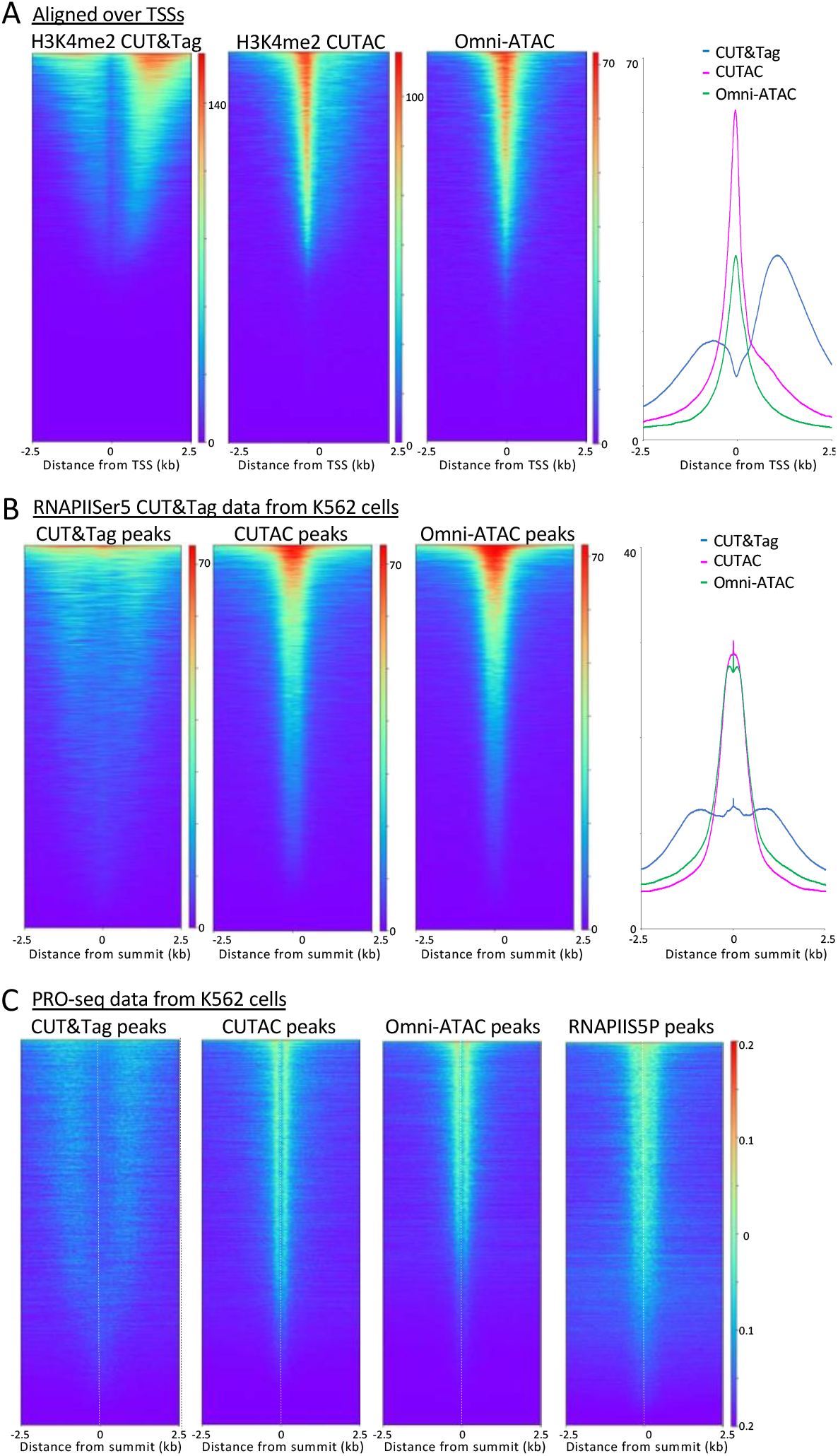
H3K4me2 CUTAC sites are coupled to transcription. (**A**) H3K4me2 fragments shift from flanking nucleosomes to the NDR upon low-salt tagmentation, corresponding closely to ATAC-seq sites. (**B**) The Serine-5 phosphate-marked initiation form of RNAPII is highly abundant over most H3K4me2 CUT&Tag, CUTAC and ATAC-seq peaks. (**C**) Run-on transcription initiates from most sites corresponding to CUTAC and ATAC-seq peaks. Both plus and minus strand PRO-seq datasets downloaded from GEO (GSM3452725) were pooled and aligned over peaks called using 3.2 million fragments sampled from H3K4me2 CUT&Tag, CUTAC and Omni-ATAC datasets, and also from pooled CUT&Tag replicate datasets for K562 RNA Polymerase II Serine-5 phosphate.

To determine whether CUTAC sites are also sites of transcription initiation in general, we aligned CUT&Tag RNA Polymerase II (RNAPII) Serine-5 phosphate (RNAPIIS5P) CUT&Tag data over H3K4me2 CUT&Tag and CUTAC and Omni-ATAC peaks ordered by RNAIIS5P peak intensity. When displayed as heatmaps or average plots, CUTAC datasets show a conspicuous shift into the NDR from flanking nucleosomes (**Figure 6B**).

Mammalian transcription also initiates at many enhancers, as shown by transcriptional run-on sequencing, which identifies sites of RNAPII pausing whether or not a stable RNA product is normally produced (28). Accordingly, we aligned RNAPII-profiling PRO-seq data for K562 cells over H3K4me2 CUT&Tag and CUTAC and Omni-ATAC sites, displayed as heatmaps and ordered by PRO-Seq signal intensity. The CUT&Tag sites showed broad enrichment of PRO-seq signals offset ~1 kb on either side, whereas PRO-seq signals were tightly centered around CUTAC sites, with similar results for Omni-ATAC sites (**Figure 6C**). Interestingly, alignment around TSSs, RNAPolIIS5P or PRO-seq data resolved immediately flanking H3K4me2-marked nucleosomes in CUT&Tag data, which is not seen for the same data aligned on signal midpoints (**Figs. 3, 5**). Such alignment of+1 and −1 nucleosomes next to fixed NDR boundaries is consistent with nucleosome positioning based on steric exclusion (29). Furthermore, the split in PRO-seq occupancies around NDRs defined by CUTAC and Omni-ATAC implies that the steady-state location of most engaged RNAPII is immediately downstream of the NDR from which it initiated. About 80% of the CUTAC sites showed enrichment of PRO-Seq signal downstream, confirming that the large majority of CUTAC sites correspond to NDRs representing transcription-coupled regulatory elements.

## Discussion

The correlation between sites of high chromatin accessibility and transcriptional regulatory elements, including enhancers and promoters, has driven the development of several distinct methods for genome-wide mapping of DNA accessibility for nearly two decades (30). However, the processes that are responsible for creating gaps in the nucleosome landscape are not completely understood. In part this uncertainty is attributable to variations in nucleosome positioning within a population of mammalian cells such that there is only a ~20% median difference in absolute DNA accessibility between DNaseI hypersensitive sites and non-hypersensitive sites genome-wide (12). This suggests that DNA accessibility is not the primary determinant of gene regulation, and contradicts the popular characterization of accessible DNA sites as “open” and the lack of accessibility as “closed”. Moreover, there are multiple dynamic processes that can result in nucleosome depletion, including transcription, nucleosome remodeling, transcription factor binding, and replication, so that the identification of a presumed regulatory element by chromatin accessibility mapping leaves open the question as to how accessibility is established and maintained. Our CUTAC mapping method now provides a physical link between a transcription-coupled process and DNA hyperaccessibility by showing that anchoring of Tn5 to a nucleosome mark laid down by transcriptional events immediately downstream identifies presumed gene regulatory elements that are indistinguishable from those identified by ATAC-seq. The equivalence of CUTAC and ATAC at both enhancers and promoters provides support for the hypothesis that these regulatory elements are characterized by the same regulatory architecture (31, 32).

The mechanistic basis for asserting that H3K4 methylation is a transcription-coupled event is well-established (20, 21). In all eukaryotes, H3K4 methylation is catalyzed by COMPASS/SET1 and related enzyme complexes, which associate with the C-terminal domain (CTD) of the large subunit of RNAPII when Serine-5 of the tandemly repetitive heptad repeat of the CTD is phosphorylated following transcription initiation. The enrichment of dimethylated and trimethylated forms of H3K4 is thought to be the result of exposure of the H3 tail to SET1/MLL during RNAPII stalling just downstream of the TSS, so that these modifications are coupled to the onset of transcription (21). Therefore, our demonstration that Tn5 tethered to H3K4me2 or H3K4me3 histone tail residues efficiently tagments accessible sites, implies that accessibility at regulatory elements is created by events immediately following transcription initiation. This mechanistic interpretation is supported by the mapping of CUTAC sites just upstream of RNAPII, and is consistent with the recent demonstration that PRO-seq data can be used to accurately impute “active” histone modifications (16). Thus CUTAC identifies active promoters and enhancers that produce enhancer RNAs, which might help explain why ~95% of ATAC-seq peaks are detected by CUTAC and vice-versa (**Figure 5B-C**).

CUTAC also provides practical advantages over other chromatin accessibility mapping methods. Like CUT&Tag-direct, all steps from frozen nuclei to purified sequencing-ready libraries for the data presented here were performed in a day in single PCR tubes on a home workbench. As it requires only a simple modification of one step in the CUT&Tag protocol, CUTAC can be performed in parallel with an H3K4me2 CUT&Tag positive control and other antibodies using multiple aliquots from each population of cells to be profiled. We have shown that three distinct protocol modifications, dilution, removal and post-wash tagmentation yield high-quality results, providing flexibility that might be important for adapting CUTAC to nuclei from diverse cell types and tissues.

Although a CUT&Tag-direct experiment requires a day to perform, and ATAC-seq can be performed in a few hours, this disadvantage of CUTAC is offset by the better control of data quality with CUTAC as is evident from the large variation in ATAC-seq data quality between laboratories (**Table 1**). In contrast, CUT&Tag is highly reproducible using native or lightly cross-linked cells or nuclei (19), and as shown here H3K4me2 CUTAC maps regulatory elements with sensitivity and signal-to-noise comparable to the best ATAC-seq datasets, even better when larger fragments are computationally excluded. Although H3K4me2 CUT&Tag datasets have lower background than CUTAC datasets run in parallel, the combination of the two provides both highest data quality (CUT&Tag) and precise mapping (CUTAC) using the same H3K4me2 antibody. Therefore, we anticipate that current CUT&Tag users and others will find the CUTAC option to be an attractive alternative to other DNA accessibility mapping methods for identifying transcription-coupled regulatory elements.

## Materials and Methods

### Biological materials

Human K562 cells were purchased from ATCC (CCL-243) and cultured following the supplier’s protocol. H1 ES cells were obtained from WiCell (WA01-lot#WB35186) and cultured following NIH 4D Nucleome guidelines (https://data.4dnucleome.org/protocols/50f8300d-400f-4ce1-8163-42f417cbbada/). We used the following antibodies: Guinea Pig anti-Rabbit IgG (Heavy& Light Chain) antibody (Antibodies-Online ABIN101961 or Novus NBP1-72763), Rabbit anti-mouse (Abcam ab46540), H3K4me1 (Epicypher 13-0026, lot 28344001), H3K4me2 (Epicypher 13-0027 and Millipore 07-030, lot 3229364), H3K4me3 (Active Motif, 39159), H3K9me3 (Abcam ab8898, lot GR3302452-1), H3K27me3 (Cell Signaling Technology, 9733, Lot 14), H3K27ac (Millipore, MABE647), H3K36me3 (Epicypher #13-0031, lot 18344001) and NPAT (Thermo Fisher Scientific, PA5-66839). The pAG-Tn5 fusion protein used in many of these experiments was a gift from Epicypher, Inc. (#15-1117 lot #20142001-C1).

### CUT&Tag-direct and CUTAC

Log-phase human K562 or H1 embryonic stem cells were harvested and prepared for nuclei in a hypotonic buffer with 0.1% Triton-X100 essentially as described (33). A detailed, step-by-step nuclei preparation protocol can be found at https://www.protocols.io/view/bench-top-cut-amp-tag-bcuhiwt6.

CUT&Tag-direct was performed as described (19), except that all CUTAC experiments were done on a home laundry room counter (**Figure 1–figure supplement 1**) with 32 samples run in parallel mostly over the course of a single ~8 hour day. A detailed step-by-step protocol including the three CUTAC options used in this study can be found at https://www.protocols.io/view/cut-amp-tag-direct-with-cutac-bmbfk2jn. Except as noted, all experiments were performed on a workbench in a home laundry room (**Figure S1**) Briefly, nuclei were thawed, mixed with activated Concanavalin A beads and magnetized to remove the liquid with a pipettor and resuspended in Wash buffer (20 mM HEPES pH 7.5, 150 mM NaCl, 0.5 mM spermidine and Roche EDTA-free protease inhibitor). After successive incubations with primary antibody (1-2 hr) and secondary antibody (0.5-1 hr) in Wash buffer, the beads were washed and resuspended in pA(G)-Tn5 at 12.5 nM in 300- Wash buffer (Wash buffer containing 300 mM NaCl) for 1 hr. Incubations were performed at room temperature either in bulk or in volumes of 25-50 μL in low-retention PCR tubes. For CUT&Tag, tagmentation was performed for 1 hr in 300-Wash buffer supplemented with 10 mM MgCl2 in a 50 μL volume. For CUTAC, tagmentation was performed in low-salt buffer with varying components, volumes and temperatures as described for each experiment in the figure legends. In “dilution” tagmentation, tubes containing 25 μL of pA(G)-Tn5 incubation solution and 2 mM or 5 mM MgCl2 solutions were preheated to 37°C. Tagmentation solution (475 μL) was rapidly added to the tubes and incubated for times and temperatures as indicated. In “removal” tagmentation, tubes were magnetized, liquid was removed, and 50 μL of ice-cold 10 mM TAPS, 5 mM MgCl2 was added, followed by incubation for times and temperatures as indicated. The “postwash” protocol is identical to the CUT&Tag-direct protocol except that tagmentation was performed in 10 mM TAPS, 5 mM MgCl2 at 37°C as indicated. In “add-back” tagmentation, the post-wash protocol was used with 10 mM TAPS, 5 mM MgCl2 supplemented with pA(G)-Tn5 and incubated at 37°C as indicated.

Following tagmentation, CUT&Tag and CUTAC samples were chilled and magnetized, liquid was removed, and beads were washed in 50 μL 10 mM TAPS pH8.5, 0.2 mM EDTA then resuspended in 5 μL 0.1% SDS, 10 μL TAPS pH8.5. Following incubation at 58°C, SDS was neutralized with 15 μL of 0.67% Triton-X100, and 2 μL of 10 mM indexed P5 and P7 primer solutions were added. Tubes were chilled and 25 μL of NEBNext 2x Master mix was added and vortexed. Gap-filling and 12 cycles of PCR were performed using an MJ PTC-200 Thermocycler. Clean-up was performed by addition of 65 μL SPRI bead slurry following manufacturer’s instructions, eluted with 20 μL 1 mM Tris-HCl pH 8, 0.1 mM EDTA and 2 μL was used for Agilent 4200 Tapestation analysis. The barcoded libraries were mixed to achieve equimolar representation as desired aiming for a final concentration as recommended by the manufacturer for sequencing on an Illumina HiSeq 2500 2-lane Turbo flow cell.

### Data processing and analysis

Paired-end reads were aligned to hg19 using Bowtie2 version 2.3.4.3 with options: ╌ end-to-end -very-sensitive ╌no-unal ╌no-mixed ╌no-discordant ╌phred33 -I 10 - X 700. Tracks were made as bedgraph files of normalized counts, which are the fraction of total counts at each basepair scaled by the size of the hg19 genome. Peaks were called using MACS2 version 2.2.6 callpeak -f BEDPE -g hs -p le-5 -keep-dup all -SPMR. Heatmaps were produced using deepTools 3.3.1.

To produce the scatterplot (**Figure 4–figure supplement 1**) and correlation matrix (**Figure 4E**), we first removed fragments overlapping any repeat-masked region in hg19, then sampled 3.2 million fragments from each of the 11 datasets and called peaks on the merged data using MACS2. As previously described (34), we used a CUTAC IgG negative control, summing normalized counts within peaks and removing peaks above a threshold of the 99^th^ percentile of normalized count sums (46,561 final peaks).

A detailed step-by-step Data Processing and Analysis Tutorial can be found at https://www.protocols.io/view/cut-amp-tag-data-processing-and-analysis-tutorial-bjk2kkye.

## Acknowledgments

We thank Terri Bryson, Christine Codomo for sample processing, the Fred Hutch Genomics Shared Resource for DNA sequencing, members of our laboratory for helpful discussions and Paul Talbert for critically reading the manuscript. S. H. is an Investigator of the Howard Hughes Medical Institute. This work was supported by the Howard Hughes Medical Institute (S.H.), grants R01 HG010492 (S.H.) and R01 GM108699 (K.A.) from the National Institutes of Health, and an HCA Seed Network grant from the Chan-Zuckerberg Initiative (S.H.).

## Supplementary Files

**Supplementary File 1.** MSExcel spreadsheets of metadata information for each figure panel and track (Tab 1), for each dataset in GEO (Tab 2), and for other GEO/SRA database files (Tab 3) used in the study.

### Supplementary figures

**Figure 1-figure supplement 1.**
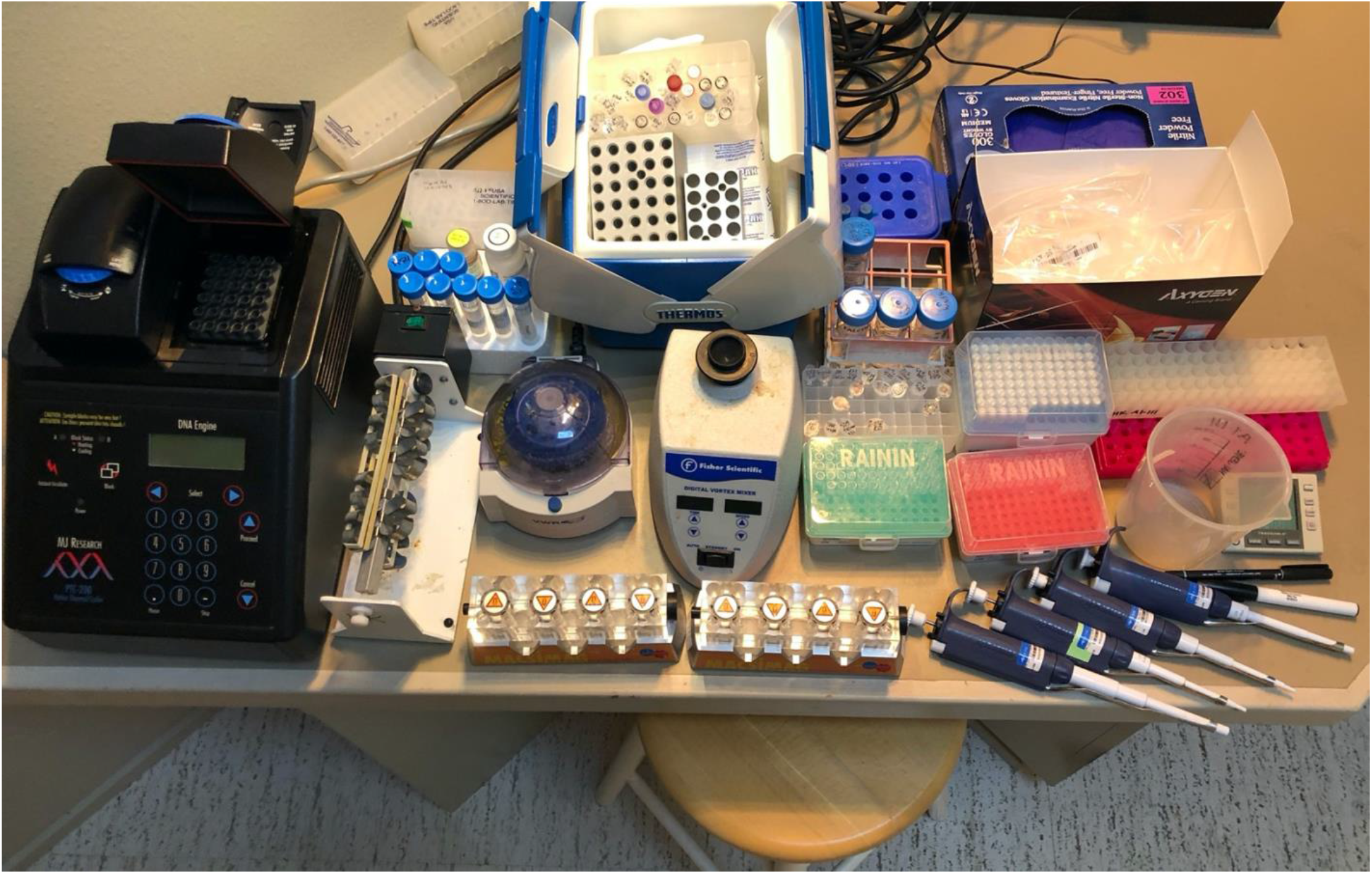
Equipment, supplies, reagents and solutions for CUT&Tag on a home workbench. All experiments starting with nuclei frozen in 10% DMSO in Mr. Frosty containers down to –80°C and held at –20°C were performed on a counter in a home laundry/utility room using stock solutions previously prepared in the lab. Following SPRI bead clean-up and liquid removal from the starting PCR tubes to fresh tubes, samples were brought into the lab for TapeStation analysis and equimolar mixing of barcoded samples followed by an SPRI clean-up of the pool and dilution for submission to the Fred Hutch Genomics Shared Resource for Illumina PE25×25 DNA sequencing. There are no hazardous materials or dangerous equipment used in the at-home protocol, however appropriate lab safety training is recommended.

**Figure 3–figure supplement 1:**
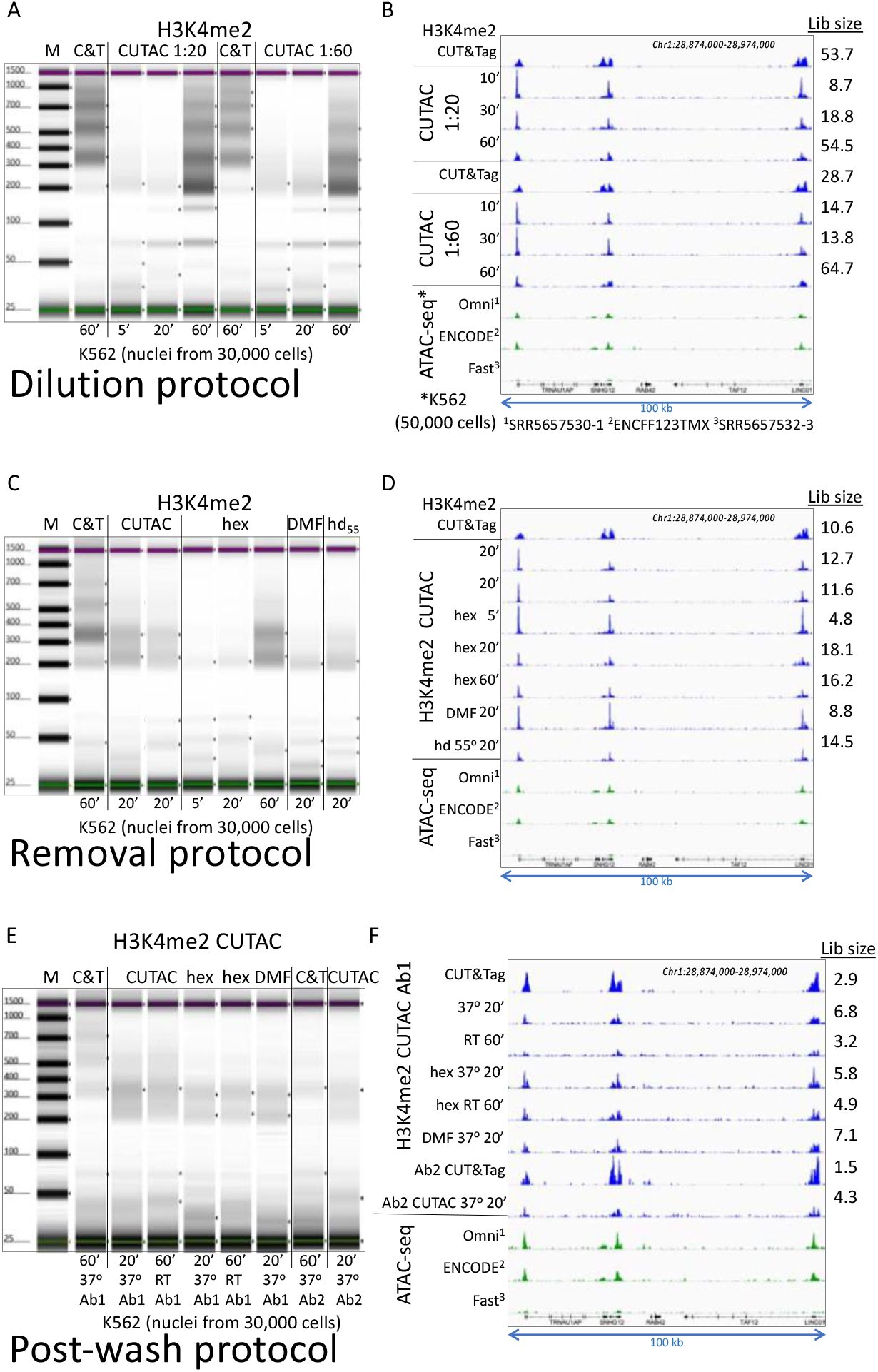
Three low-salt tagmentation protocols map chromatin hyperaccessibility: **A-B)** H3K4me2 CUT&Tag (C&T) and low-salt tagmentations using the Dilution protocol were performed at 37°C using Epicypher 13-0027 antibody and Epicypher 15-1117 pAG-Tn5 for the times indicated using pA/G-Tn5 at either 1:20 (Manufacturer’s recommendation) or 1:60 showing a bigger effect of tagmentation time than amount of pA/G-Tn5, and improvement in yield but reduction in signal-to-noise with longer tagmentations. A) Tapestation gel images showing time of tagmentation and yields based on loading 2 μL of each 20 μL sample and integrating over the 175-1000 bp range. M = markers, C&T = CUT&Tag; B) Group-autoscaled tracks showing fragment normalized count densities for sequenced libraries using 30,000 cells resolved in (A) and for published ATAC-seq data using 50,000 cells. **C-D**) Same as A-B using the Removal protocol with no additive (CUTAC), 10% 1,6-hexanediol (hex), 10% N,N-dimethylformamide (DMF) or 10% of both at 55°C (hd55) for the times indicated. All datasets were sampled down to 3.2 million and mapped to hg19. A representative 100-kb region was group-autoscaled using IGV. **E-F**) Same as A-B using the Post-wash protocol with no additive, hex and DMF at 37°C or Room temperature (21-22°C) as indicated. Ab1 is Epicypher 13-0027 and Ab2 is Millipore 07-030. Post-wash gel image was adjusted to visualize faint bands, and the lower yields are reflected in reduced Estimated library size (Lib size, millions of fragments), calculated by the Mark Duplicates program in Picard tools.

**Figure 3–figure supplement 2:**
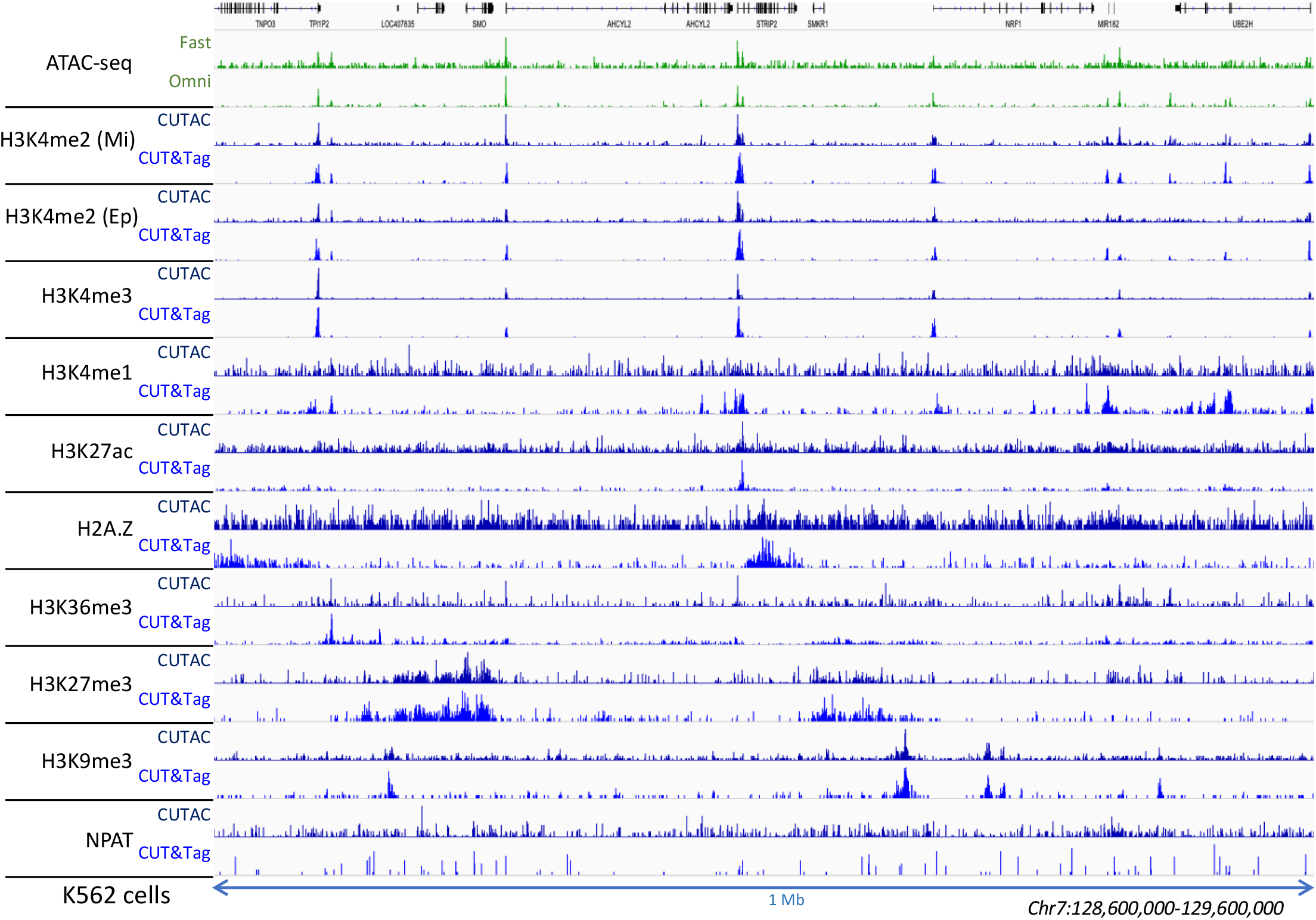
Low-salt tagmentation using various antibodies. Two H3K4me2 antibodies were used: Millipore 07-030 lot 3229364 (Mi) and Epicypher 13-0027 (Ep) and provided similar results. CUTAC was done using the Removal protocol and incubated 10 min 37°C.

**Figure 3–figure supplement 3:**
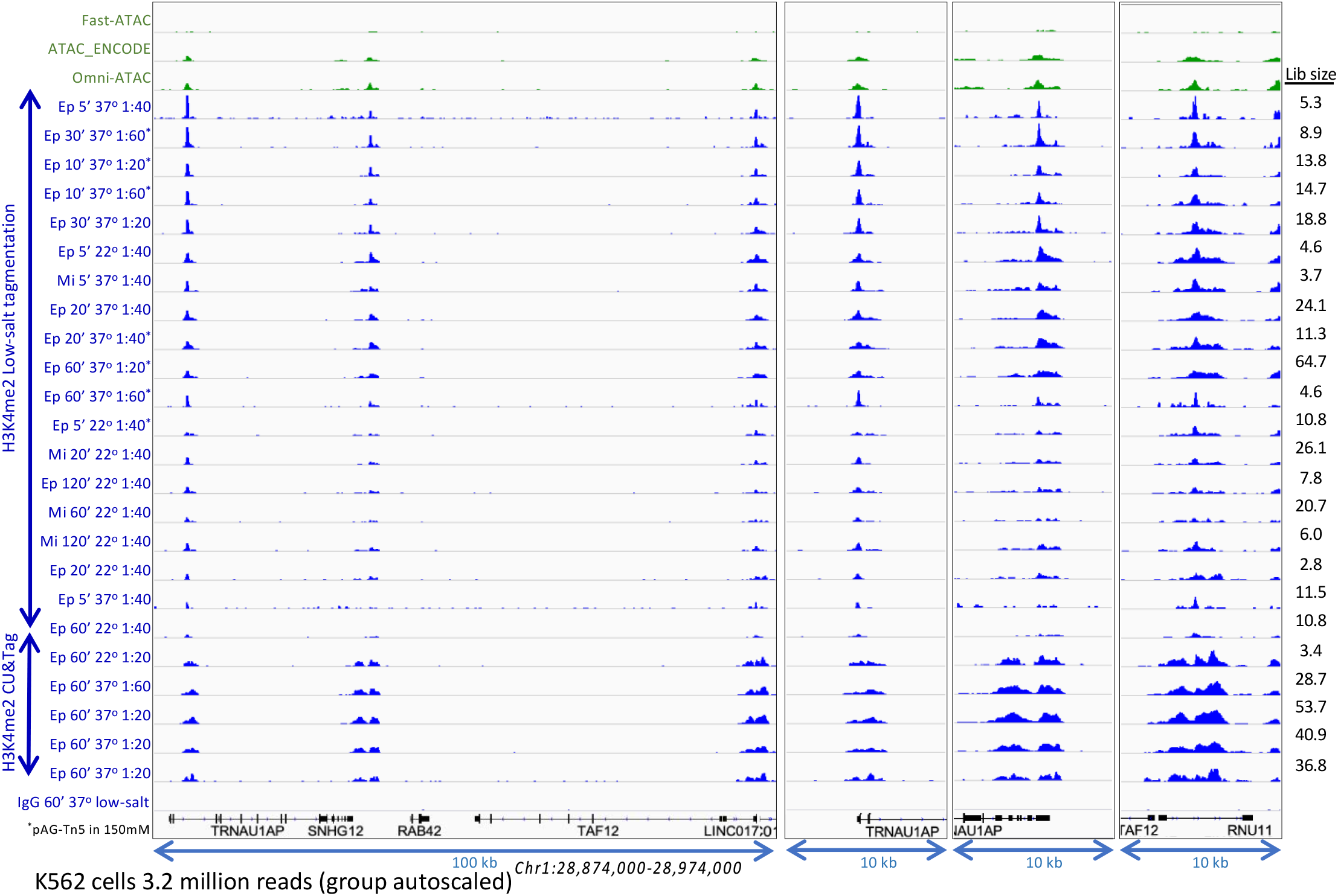
Optimization of low-salt tagmentation conditions: H3K4me2 CUT&Tag and low-salt tagmentation were performed using either a rabbit polyclonal [Millipore 07-030 lot 3229364 (Mi)] or rabbit monoclonal [Epicypher 13-0027 (Ep)] antibody with pAG-Tn5 (Epicypher 15-1117 lot #20142001-C1) at the indicated dilutions. Dilution tagmentation in 2 mM MgCl2 was used at either 22°C or 37°C. Raw paired-end reads were mapped to hg19. A representative 100-kb region is shown (left) and expanded (right) around active promoters and group-autoscaled using IGV. Estimated library size (Lib size) was calculated by the Mark Duplicates program in Picard tools.

**Figure 3–figure supplement 4:**
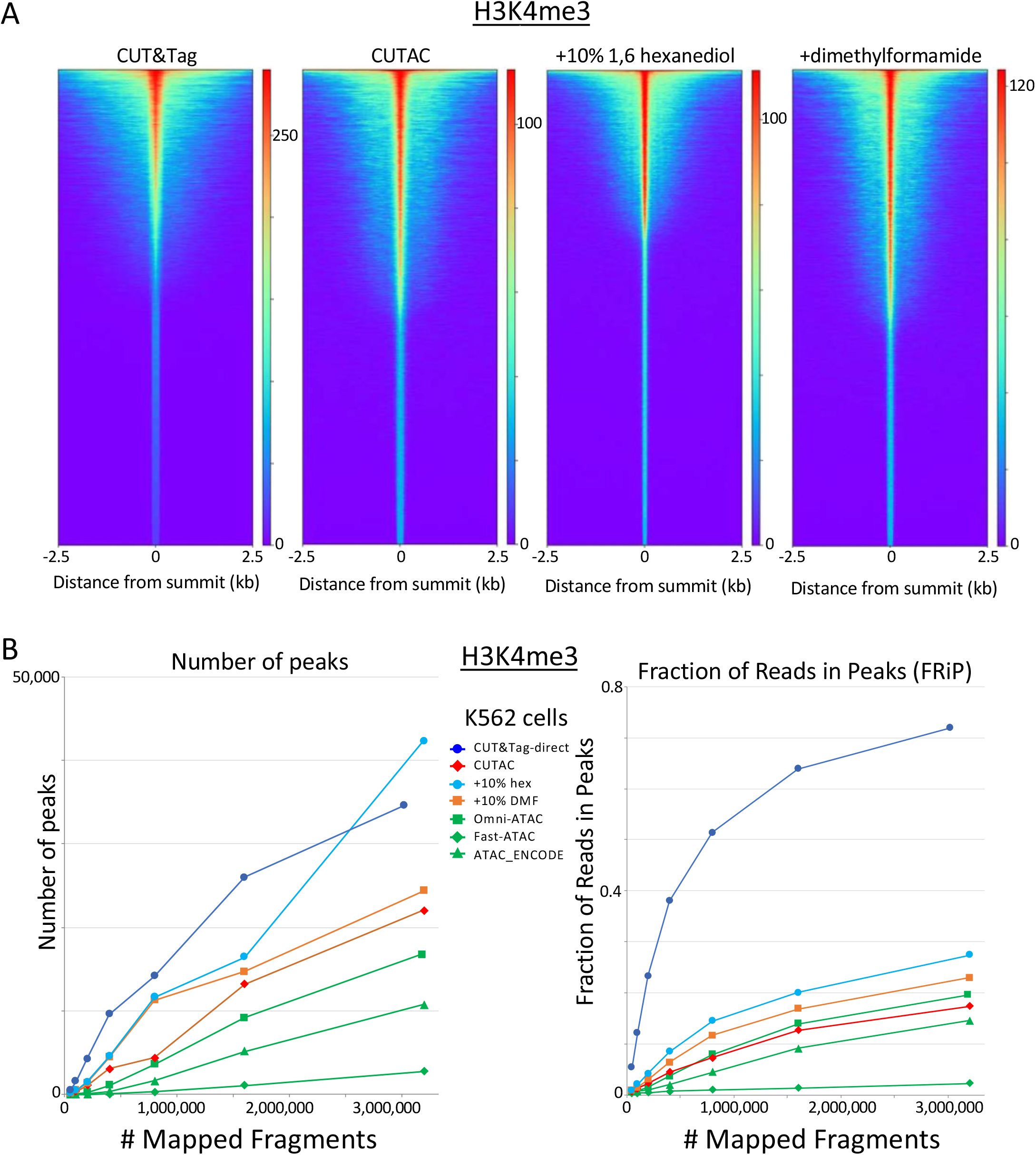
H3K4me3 CUTAC shows peak narrowing and improved peak-calling with addition of 1,6-hexanediol. (**A**) Peak narrowing is observed with addition of 10% 1,6-hexanediol although not with N,N-dimethylformamide. (**B**) Addition of 10% 1,6-hexanediol increases the number of peaks called and improves FRiP, with possible slight improvements seen with addition of N,N-dimethylformamide.

**Figure 4-figure supplement 1:**
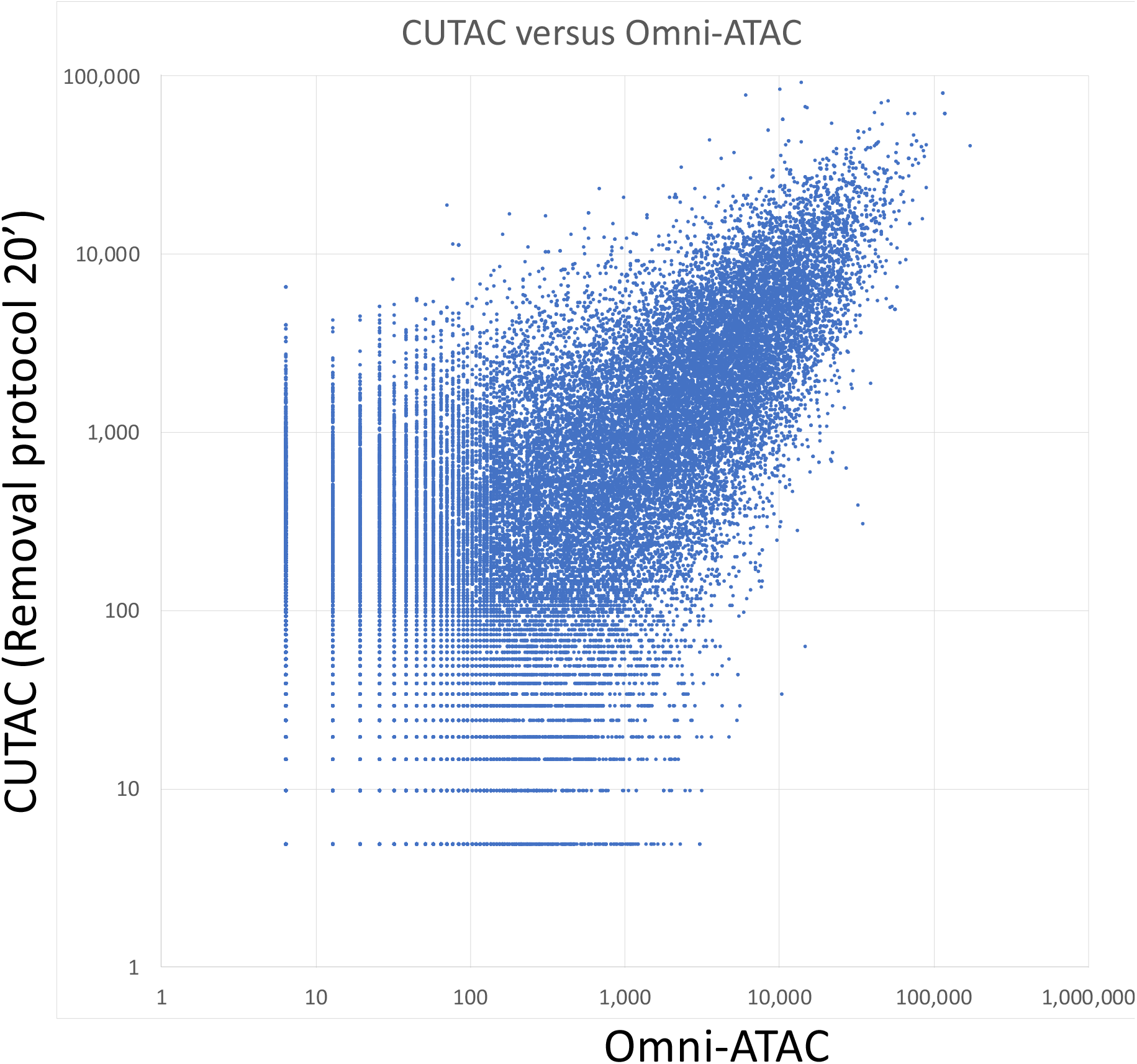
Log_10_ scatterplot of CUTAC versus Omni-ATAC (R^2^= 0.53). Samples of 3.2 million fragments were pooled and MACS2 was used to call peaks. The sums of normalized counts spanning each peak for the two datasets were plotted.

**Figure 5–figure supplement 1:**
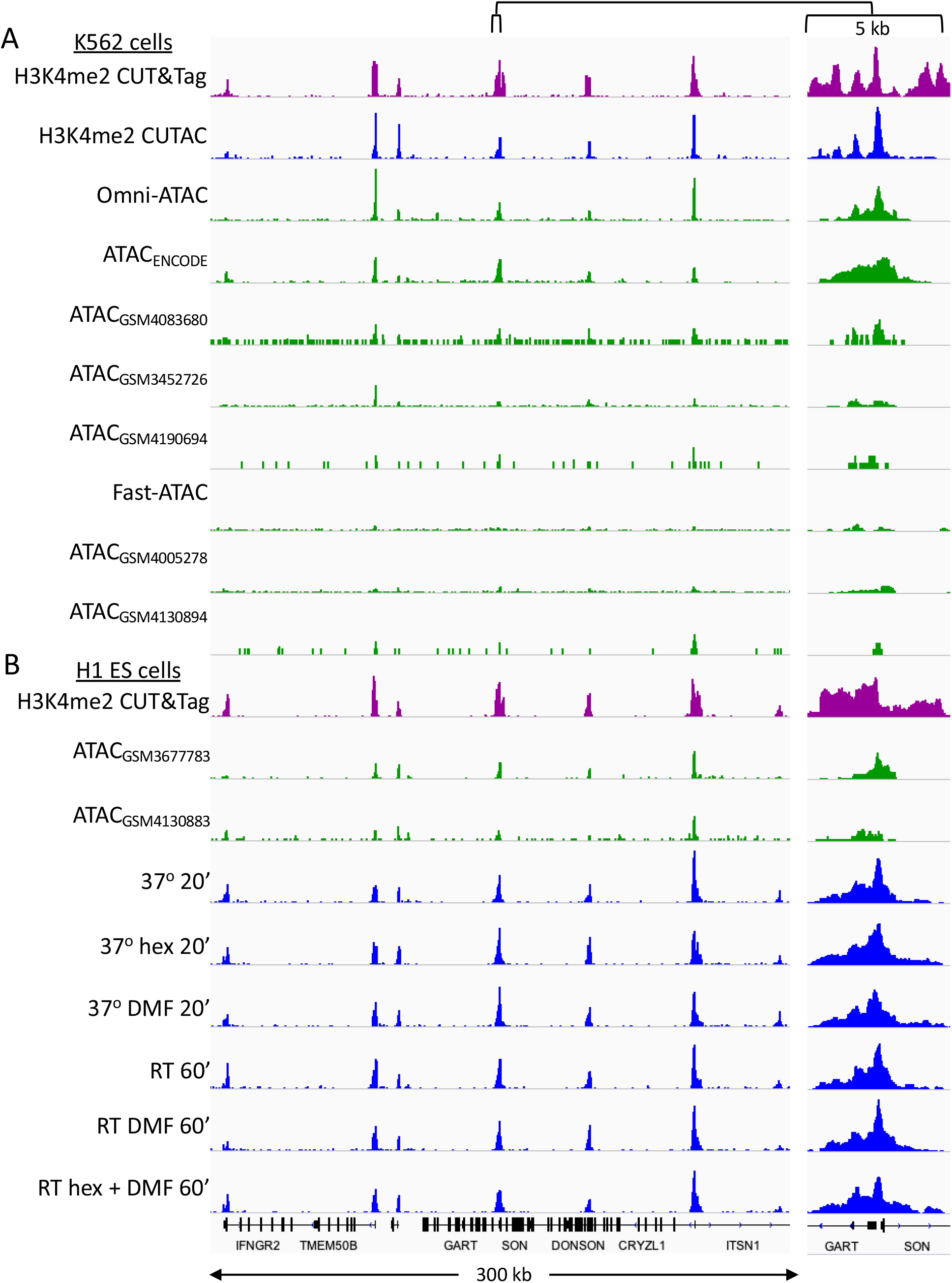
CUTAC data quality is similar to that of the best ATAC-seq datasets. Tracks over a representative region for K56^n^ datasets listed in Supplementary Table 1. Samples are ordered by decreasing FRiP. (**A**) K562 cells (**B**) H1 ES cells (Post-wash protocol).

**Figure 5–figure supplement 2:**
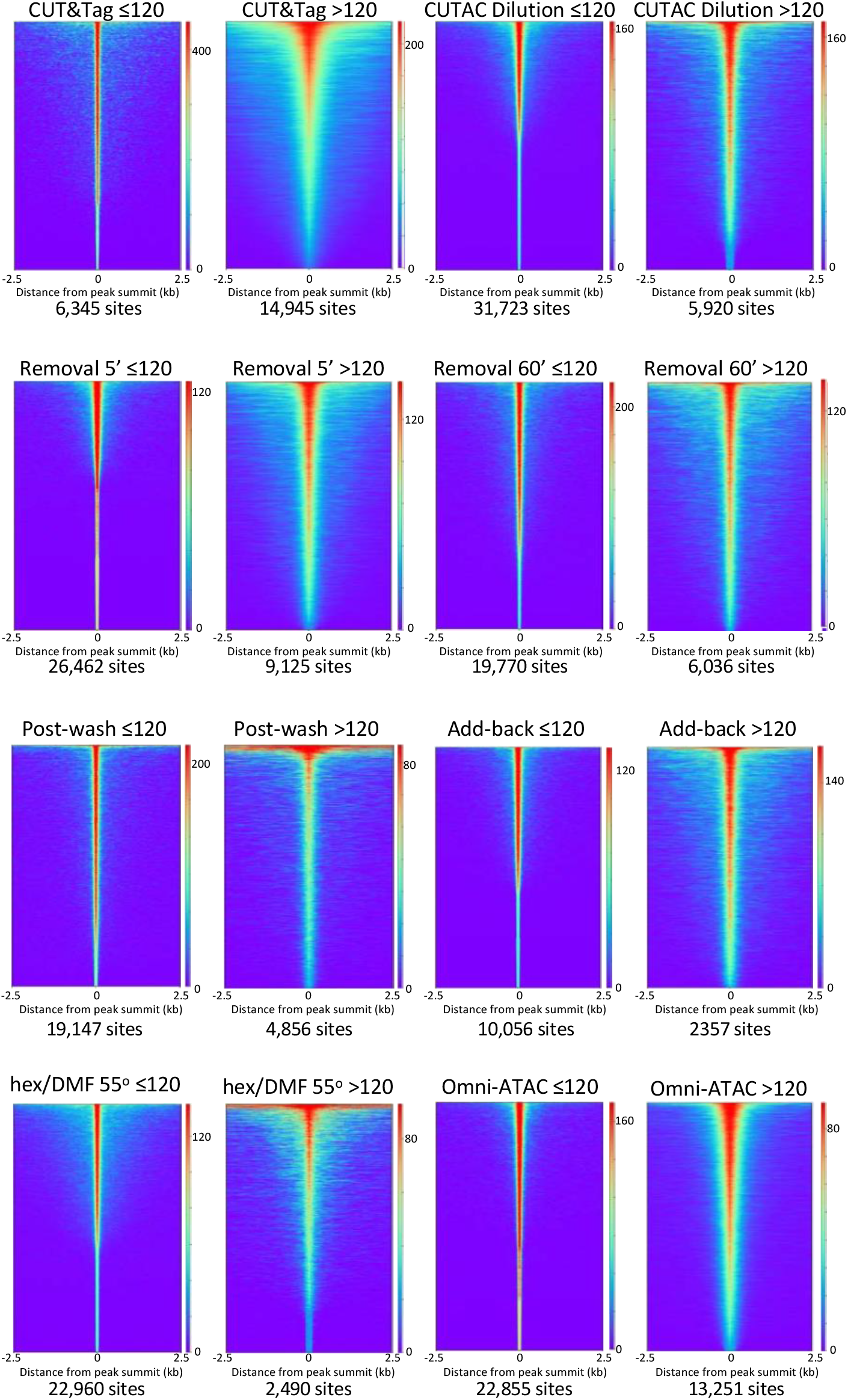
Smaller fragments (≤120 bp) dominate NDRs. See Figure 2E. Additional comparisons of small (≤120 bp) and large (>120) fragments from diverse H3K4me2 CUTAC datasets used in this study show consistent narrowing for small fragments around their summits. For each dataset a 3.2 million fragment random sample was split into small and large fragment groups, MAC2 was used to call peaks and heatmaps were ordered by density over the peak midpoints (sites).

**Figure 5–figure supplement 3:**
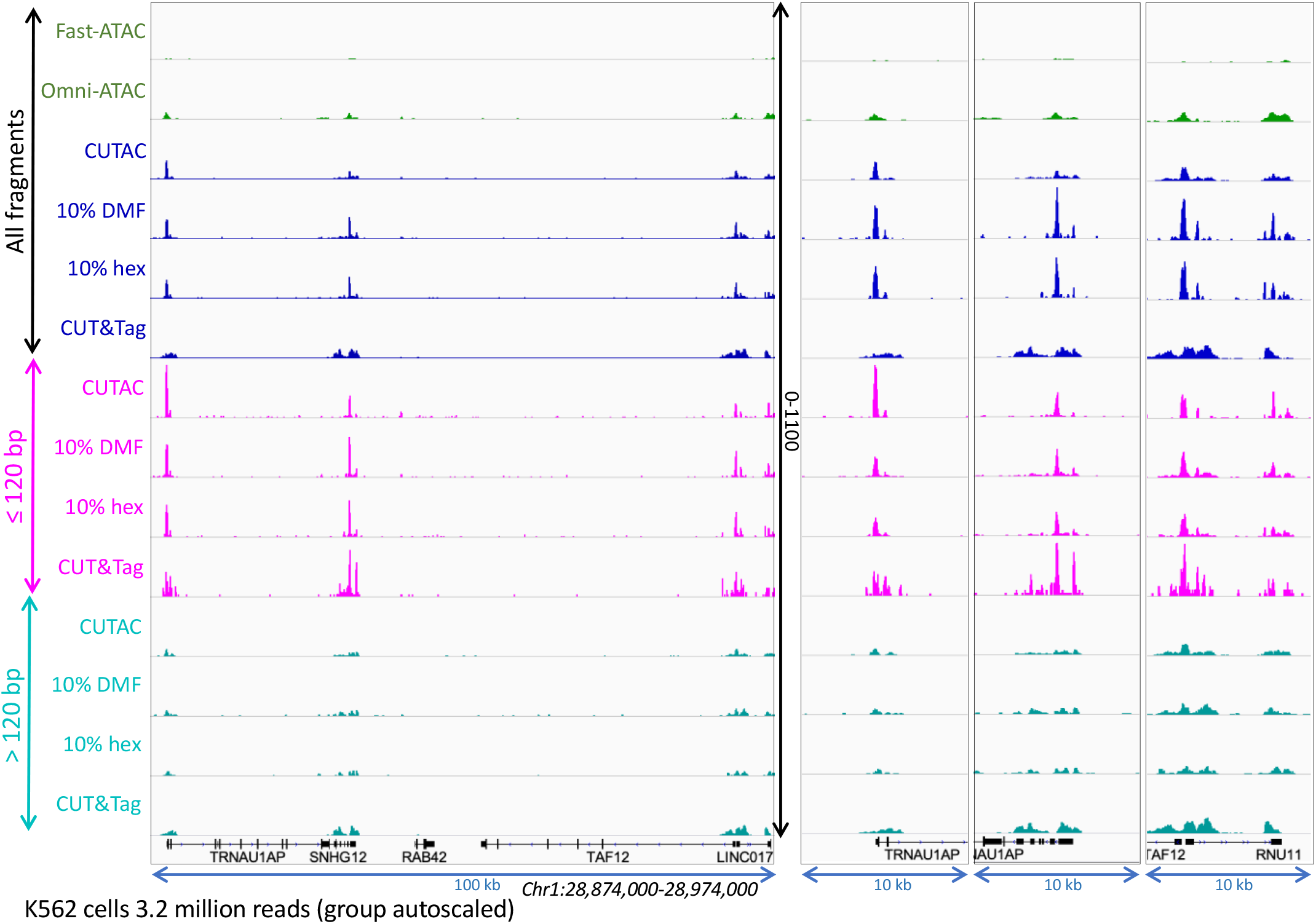
Small CUTAC fragments improve peak resolution. The representative region shown in Figs. S3-S4 is shown for all 3.2 million fragments and for ≤120 bp and >120 bp groups.

**Figure 6-figure supplement 1:**
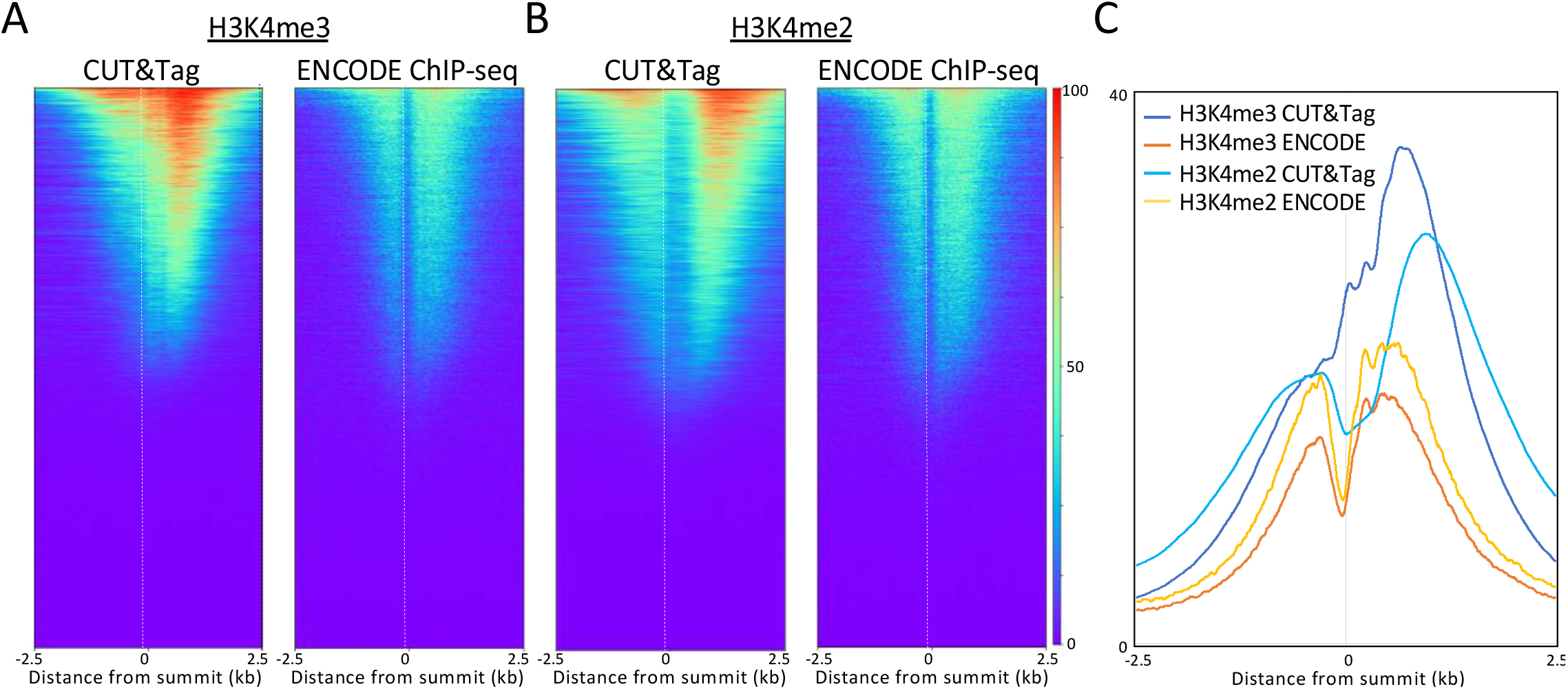
ChlP-seq confirms CUT&Tag localization for H3K4me2/3 flanking active promoters. (**A**) Comparison of H3K4me3 CUT&Tag to ENCODE ChlP-seq. (**B**) Same as (A) for H3K4me2. (**C**) CUT&Tag shows that H3K4me2 is centered farther from the NDR than is H3K4me3.

## Notes

https://www.protocols.io/view/cut-amp-tag-direct-with-cutac-bmbfk2jn

### Competing Interest Statement

S.H. and H.S.K. have filed patent applications on related work.

### Summary of Updates

This version further updates CUT&Tag@home with a description of the CUTAC method, and includes an updated link to our Protocols.io site with a step-by-step procedure for performing CUTAC as part of a CUT&Tag-direct experiment.

## References

1. Weintraub H, Groudine M. Chromosomal subunits in active genes have an altered conformation. Science. 1976;193:848–56.

2. Reeves R. Nucleosome structure of Xenopus oocyte amplified ribosomal genes. Biochemistry (Mosc). 1978;17(23):4908–16.

3. Jack RS, Eggert H. Restriction enzymes have limited access to DNA sequences in Drosophila chromosomes. EMBO J. 1990;9(8):2603–9.

4. Gottschling DE. Telomere-proximal DNA in Saccharomyces cerevisiae is refractory to methyltransferase activity in vivo. Proceedings of the National Academy of Sciences of the United States of America. 1992;89:4062–5.

5. Schwartz YB, Kahn TG, Pirrotta V. Characteristic low density and shear sensitivity of cross-linked chromatin containing polycomb complexes. Mol Cell Biol. 2005;25(1):432–9.

6. Bownes M. Preferential insertion of P elements into genes expressed in the germ-line of Drosophila melanogaster. Mol Gen Genet. 1990;222(2–3):457#x2013;60.

7. Crawford GE, Holt IE, Mullikin JC, Tai D, Blakesley R, Bouffard G, et al. Identifying gene regulatory elements by genome-wide recovery of DNase hypersensitive sites. Proc Natl Acad Sci U S A. 2004;101(4):992–7.

8. Dorschner MO, Hawrylycz M, Humbert R, Wallace JC, Shafer A, Kawamoto J, et al. High-throughput localization of functional elements by quantitative chromatin profiling. Nat Methods. 2004;1(3):219–25.

9. Giresi PG, Kim J, McDaniell RM, Iyer VR, Lieb JD. FAIRE (Formaldehyde-Assisted Isolation of Regulatory Elements) isolates active regulatory elements from human chromatin. Genome Res. 2007;17(6):877–85.

10. Auerbach RK, Euskirchen G, Rozowsky J, Lamarre-Vincent N, Moqtaderi Z, Lefrancois P, et al. Mapping accessible chromatin regions using Sono-Seq. Proc Natl Acad Sci USA. 2009;106(35):14926–31.

11. Buenrostro JD, Giresi PG, Zaba LC, Chang HY, Greenleaf WJ. Transposition of native chromatin for fast and sensitive epigenomic profiling of open chromatin, DNA-binding proteins and nucleosome position. Nat Methods. 2013;10(12):1213–8.

12. Chereji RV, Eriksson PR, Ocampo J, Prajapati HK, Clark DJ. Accessibility of promoter DNA is not the primary determinant of chromatin-mediated gene regulation. Genome Res. 2019;29(12):1985–95.

13. Oberbeckmann E, Wolff M, Krietenstein N, Heron M, Ellins JL, Schmid A, et al. Absolute nucleosome occupancy map for the Saccharomyces cerevisiae genome. Genome Res. 2019;29(12):1996–2009.

14. Karabacak Calviello A, Hirsekorn A, Wurmus R, Yusuf D, Ohler U. Reproducible inference of transcription factor footprints in ATAC-seq and DNase-seq datasets using protocol-specific bias modeling. Genome Biol. 2019;20(1):42.

15. Kornberg RD, Lorch Y. Primary Role of the Nucleosome. Mol Cell. 2020;79(3):371–5.

16. Wang Z, Chivu AG, Choate LA, Rice EJ, Miller DC, Chu T, et al. Accurate imputation of histone modifications using transcription. biorxiv. 2020;https://doi.org/10.1101/2020.04.08.032730.

17. Schmid M, Durussel T, Laemmli UK. ChIC and ChEC; genomic mapping of chromatin proteins. Mol Cell. 2004;16(1):147–57.

18. Kaya-Okur HS, Wu SJ, Codomo CA, Pledger ES, Bryson TD, Henikoff JG, et al. CUT & Tag for efficient epigenomic profiling of small samples and single cells. Nat Commun. 2019;10:1930.

19. Kaya-Okur HS, Janssens DH, Henikoff JG, Ahmad K, Henikoff S. Efficient low-cost chromatin profiling with CUT & Tag. Nature Protocols. 2020;http://dx.doi.org/10.1038/s41596-020-0373-x.

20. Henikoff S, Shilatifard A. Histone modification: cause or cog? Trends Genet. 2011;27(10):389–96.

21. Soares LM, He PC, Chun Y, Suh H, Kim T, Buratowski S. Determinants of Histone H3K4 Methylation Patterns. Mol Cell. 2017;68(4):773–85 e6.

22. Landt SG, Marinov GK, Kundaje A, Kheradpour P, Pauli F, Batzoglou S, et al. ChIP-seq guidelines and practices of the ENCODE and modENCODE consortia. Genome Res. 2012;22(9):1813–31.

23. Corces MR, Trevino AE, Hamilton EG, Greenside PG, Sinnott-Armstrong NA, Vesuna S, et al. An improved ATAC-seq protocol reduces background and enables interrogation of frozen tissues. Nat Methods. 2017;14(10):959–62.

24. Zhang J, Lee D, Dhiman V, Jiang P, Xu J, McGillivray P, et al. An integrative ENCODE resource for cancer genomics. Nat Commun. 2020;11(1):3696.

25. Corces MR, Buenrostro JD, Wu B, Greenside PG, Chan SM, Koenig JL, et al. Lineage-specific and single-cell chromatin accessibility charts human hematopoiesis and leukemia evolution. Nat Genet. 2016;48(10):1193–203.

26. Swanson E, Lord C, Reading J, Heubeck AT, Savage AK, Green R, et al. Integrated single cell analysis of chromatin accessibility and cell surface markers. biorxiv. 2020;https://doi.org/10.1101/2020.09.04.283887.

27. Gilchrist DA, Fromm G, Dos Santos G, Pham LN, McDaniel IE, Burkholder A, et al. Regulating the regulators: the pervasive effects of Pol II pausing on stimulus-responsive gene networks. Genes Dev. 2012;26(9):933–44.

28. Kaikkonen MU, Spann NJ, Heinz S, Romanoski CE, Allison KA, Stender JD, et al. Remodeling of the enhancer landscape during macrophage activation is coupled to enhancer transcription. Mol Cell. 2013;51(3):310–25.

29. Chereji RV, Ramachandran S, Bryson TD, Henikoff S. Precise genome-wide mapping of single nucleosomes and linkers in vivo. Genome Biol. 2018;19(1):19.

30. Klein DC, Hainer SJ. Genomic methods in profiling DNA accessibility and factor localization. Chromosome Res. 2020;28(1):69–85.

31. Andersson R, Sandelin A, Danko CG. A unified architecture of transcriptional regulatory elements. Trends Genet. 2015;31(8):426–33.

32. Arnold PR, Wells AD, Li XC. Diversity and Emerging Roles of Enhancer RNA in Regulation of Gene Expression and Cell Fate. Frontiers in cell and developmental biology. 2019;7:377.

33. Skene PJ, Henikoff S. An efficient targeted nuclease strategy for high-resolution mapping of DNA binding sites. eLife. 2017;6:e21856.

34. Meers MP, Bryson TD, Henikoff JG, Henikoff S. Improved CUT & RUN chromatin profiling tools. eLife. 2019;8:e46314.

35. Moore JE, Purcaro MJ, Pratt HE, Epstein CB, Shoresh N, Adrian J, et al. Expanded encyclopaedias of DNA elements in the human and mouse genomes. Nature. 2020;583(7818):699–710.

